# RAP2.6 negatively regulates Arabidopsis submergence tolerance through GSNOR1-dependent redox and hormonal signaling

**DOI:** 10.64898/2026.10.25.761742

**Authors:** Ashim Kumar Das, Mohammad Golam Mostofa, Da-Sol Lee, Raisul Awal Mahmood, Rafiqul Alam, Adil Hussain, Nusrat Jahan Methela, Youn-Ji Woo, Sharmin Sultana, Chang-Wook Park, Bong-Gyu Mun, In-Jung Lee, Sunghwan Kim, Byung-Wook Yun

## Abstract

Group VII ETHYLENE RESPONSE FACTORs (ERFVIIs) are central regulators of plant responses to oxygen deprivation. However, the contributions of many other APETALA2 (AP2)/ERF transcription factors to hypoxia and flooding responses remain poorly understood. Here, we identify the group X ERF transcription factor RAP2.6 as a negative regulator of submergence tolerance in *Arabidopsis thaliana*. *RAP2.6* expression declined under hypoxia, and plants overexpressing *RAP2.6* exhibited reduced survival following dark submergence, whereas the *rap2.6-1* loss-of-function mutant showed enhanced tolerance to submergence, hypoxia, and ethanol treatment. Enhanced tolerance of *rap2.6-1* was associated with reduced membrane injury, maintenance of photosynthetic pigments and soluble sugars, and extensive metabolic remodeling. *RAP2.6* overexpression altered nitric oxide (NO), *S*-nitrosothiol (SNO), and ethylene accumulation during submergence and was associated with greater oxidative damage and enhanced abscisic acid (ABA)-responsive gene expression. Genetic analyses using *S-NITROSOGLUTATHIONE (GSNO) REDUCTASE 1* (*GSNOR1*) gain- and loss-of-function backgrounds demonstrated that GSNOR1-dependent GSNO homeostasis strongly modified RAP2.6-dependent submergence responses. Increased GSNOR1 activity suppressed the submergence-sensitive phenotype of *RAP2.6*-overexpressing plants, reduced nitro-oxidative stress, and restored carbohydrate reserves, whereas loss of GSNOR1 activity abolished the enhanced tolerance of *rap2.6-1*. GSNOR1 also modified RAP2.6-dependent expression of genes associated with oxygen sensing and ABA signaling, while its protective effect was not explained solely by changes in ethylene accumulation. Together, these findings identify RAP2.6 as a regulatory component of the low-oxygen response and reveal a functional interaction between RAP2.6 and GSNOR1-dependent redox signaling that coordinates plant hormonal, metabolic, and oxidative responses during submergence.

## Introduction

Flooding severely restricts gas diffusion into plant tissues, resulting in oxygen (O_2_) deficiency, impaired energy production, and substantial losses in plant growth and productivity (Bailey-Serres and Voesenek, 2008). Plants survive flooding through coordinated morphological, physiological, and metabolic acclimation, including escape or quiescence strategies, enhanced fermentative metabolism, redistribution of carbon reserves, and extensive hormonal reprogramming (Dichio et al., 2025; Praveen and Singh, 2026).

Among the plant hormones, ethylene is rapidly entrapped in submerged tissues and functions as an early signal that promotes acclimation to hypoxia and subsequent reoxygenation (Sasidharan et al., 2017; Hartman et al., 2019; Akter et al., 2026).

A central component of plant O_2_ sensing is the PROTEOLYSIS6 (PRT6) N-degron pathway. Under normoxia, PLANT CYSTEINE OXIDASEs (PCOs) oxidize the N-terminal cysteine of group VII ETHYLENE RESPONSE FACTORs (ERFVIIs), including RELATED TO APETALA 2.2 (RAP2.2), RAP2.3, RAP2.12, HYPOXIA-RESPONSIVE ERF 1 (HRE1), and HRE2, promoting their arginylation and proteasomal degradation. When O_2_ becomes limiting, reduced PCO activity stabilizes ERFVIIs, allowing their nuclear accumulation and activation of hypoxia-responsive genes (Giuntoli and Perata, 2017; Hartman et al., 2021; Das et al., 2026). ERFVII-dependent responses contribute to low-O_2_ adaptation not only in Arabidopsis but also in crops, including rice (*Oryza sativa*), barley (*Hordeum vulgare*), maize (*Zea mays*), and soybean (*Glycine max*) (Loreti and Perata, 2023). In addition to ERFVIIs, the APETALA2/ETHYLENE RESPONSE FACTOR (AP2/ERF) superfamily also contains many other additional transcription factors whose functions during O_2_-deprivation remain poorly characterized (Nakano et al., 2006; Dey and Corina Vlot, 2015). Several non-ERFVII factors, including ETHYLENE RESPONSIVE ELEMENT BINDING PROTEIN 1 (OsEREBP1) in rice, BnERF2.4 in rapeseed (*Brassica napus*), PhERF2 in petunia, and HvERF2.11 in barley have been linked to flooding responses, suggesting that broader AP2/ERF networks contribute to metabolic, antioxidant, and hormonal acclimation under flooding (Jisha et al., 2015; Lv et al., 2016; Yin et al., 2019; Luan et al., 2020).

Nitric oxide (NO) is another important component of low-O_2_ signaling. Cellular NO can arise through nitrate reductase (NR)-dependent and other reductive pathways, and NO-derived signals are closely connected with glutathione (GSH) and *S*-nitrosoglutathione (GSNO) metabolism. GSNO serves as a major reservoir and transnitrosating species within the cellular *S*-nitrosothiol (SNO) pool, whereas GSNO REDUCTASE1 (GSNOR1) regulates GSNO/SNO homeostasis and thereby influences NO-dependent protein modification and signaling (Das et al., 2025). NO has also been implicated in ERFVII turnover and in the phytoglobin/NO cycle that supports redox and energy metabolism under O_2_ limitation (Gibbs et al., 2015; Gupta and Igamberdiev, 2016). The precise contribution of NO to N-degron-mediated ERFVII turnover remains debated, because PCO-catalyzed N-terminal cysteine oxidation can occur without prior cysteine nitrosylation (White et al., 2017). Ethylene further intersects with this network by promoting NO depletion in submerged tissues through phytoglobin-dependent mechanisms (Hartman et al., 2019), emphasizing that NO, SNO homeostasis, and ethylene signaling are tightly coupled but not necessarily arranged in a simple linear pathway.

RAP2.6 is a group X ERF transcription factor initially characterized as a CORONATINE INSENSITIVE 1 (COI1)-dependent jasmonate-responsive factor and subsequently implicated in abscisic acid (ABA), salinity, cold, drought, and biotic stress responses (Fowler and Thomashow, 2002; Wang et al., 2008; Zhu et al., 2010; Krishnaswamy et al., 2011; Ali et al., 2013). RAP2.6 interacts with CYCLIN-DEPENDENT KINASE 8 (CDK8) and SUCROSE NONFERMENTING 1 (SNF1)-RELATED PROTEIN KINASE 2.6 (SnRK2.6), and binds GCC/DRE *cis* regulatory elements in stress-responsive genes, namely RESPONSE TO DESICCATION 29A (RD29A) and COLD-REGULATED 15A (COR15A), thereby contributing to ABA-dependent drought responses (Zhu et al., 2020). Our previous transcriptomic analyses also identified RAP2.6 among the strongly induced transcription factors following treatment with the NO donor S-nitrosocysteine (CysNO) (Hussain et al., 2016; Falak et al., 2021). These observations suggested that RAP2.6 could connect redox and hormonal signaling during O_2_ deprivation.

Here, we investigated the function of RAP2.6 during submergence using gain- and loss-of-function Arabidopsis lines together with genetic combinations affecting NO and GSNO/SNO homeostasis. We show that RAP2.6 negatively regulates tolerance to submergence, hypoxia, and ethanol and is associated with changes in carbon status, lipid/metabolite profiles, NO/SNO and ethylene accumulation, ABA-responsive signaling, and oxidative stress. Genetic analyses further demonstrate a strong functional interaction between RAP2.6 and GSNOR1-dependent GSNO/SNO homeostasis. Rather than assigning these components to a single linear pathway, our results support a model in which RAP2.6 and GSNOR1 jointly shape redox, hormonal, and metabolic outputs that determine plant survival during O_2_ limitation.

## Results

### RAP2.6 negatively regulates tolerance to submergence and hypoxia

Multiple sequence alignment of the ERFVII proteins RAP2.2, RAP2.3, RAP2.12, HRE1, and HRE2 with the group X ERFs RAP2.6 and RAP2.6L confirmed conservation of the AP2 DNA-binding domain (Fig. S1). Publicly available Arabidopsis eFP Browser data further indicated that RAP2.6 transcript abundance decreases under hypoxia, in contrast to the induction of several ERFVII genes and RAP2.6L (Fig. S2). To determine whether RAP2.6 contributes to low-O_2_ tolerance, we compared the responses of Col-0 plants, *RAP2.6*-overexpressing plants (*35S::RAP2.6*), and the *rap2.6-1* loss-of-function mutant plants under 5 d of dark submergence followed by 7 d of recovery. *RAP2.6*-overexpressing plants exhibited lower survival than Col-0, whereas *rap2.6-1* mutant plants showed enhanced survival (Fig. 1A, B). The *rap2.6-1* mutant also displayed lower electrolyte leakage and maintained higher photosynthetic pigment and total soluble sugar levels after submergence, whereas the opposite trend was observed in *35S::RAP2.6* plants (Fig. 1C-H). Consistent with the whole-plant phenotype, *35S::RAP2.6* seedlings showed lower green-cotyledon percentages after hypoxia or ethanol treatment, while *rap2.6-1* seedlings were more tolerant (Fig. 1I-L). These results identify RAP2.6 as a negative regulator of responses to O_2_ limitation.

**Fig. 1.**
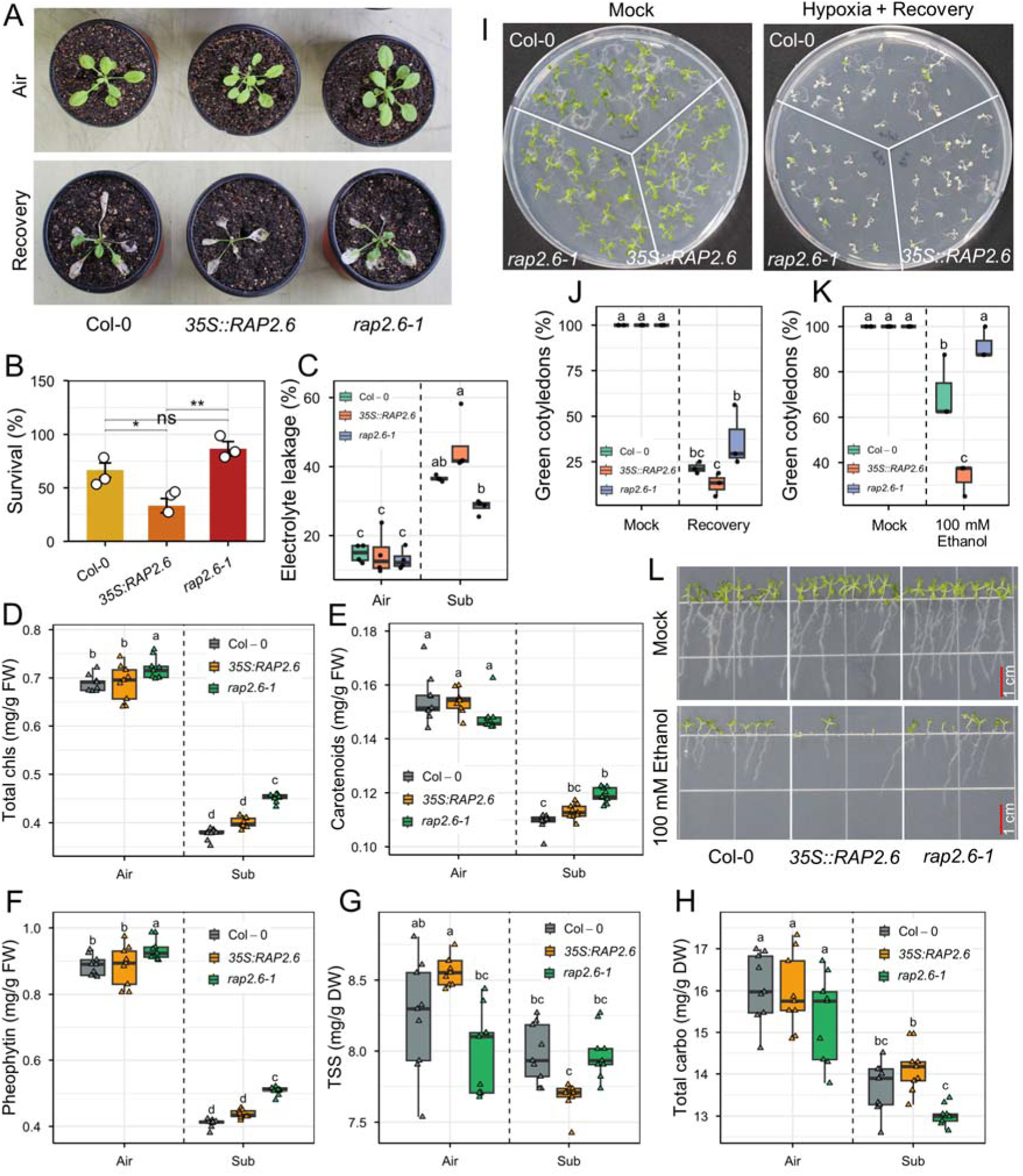
Evaluation of submergence tolerance in RAP2.6. (A) Phenotypes of Col-0, *35S::RAP2.6*, and *rap2.6-1* plants were photographed before submergence (air) and after 5 days of dark submergence, followed by 7 days of recovery. (B) Survival rates were calculated after 7 days of recovery. Bars represent means ± SE (*n* = 3) of three independent experiments (for one experiment, >10 plants were scored for each genotype). (C) Electrolyte leakage, (D) total chlorophyll (chls), (E) carotenoids, (F) pheophytin, (G) total soluble sugar (TSS), and (H) total carbohydrate (carbo) in Col-0, *35S::RAP2.6*, and *rap2.6-1* plants exposed to air or 2 days of dark submergence (means ± SE, *n* = 4 biological replicates for EL, *n* = 9 technical replicates from 3 biological replicates). (I) Phenotypes of Col-0, *35S::RAP2.6*, and *rap2.6-1* seedlings grown on ½ MS media for 7 days, followed by 1 and 3 days of hypoxia and recovery, respectively. (J) Green cotyledon percentage was calculated after 3 days of recovery. Bars represent means ± SE (*n* = 3) of three independent experiments (for one experiment, >14 plants were scored for each genotype). (K) Phenotypes and (L) green cotyledon percentage of Col-0, *35S::RAP2.6*, and *rap2.6-1* seedlings grown on ½ MS media for 10 days, with or without 100 mM Ethanol. Bars represent means ± SE (*n* = 3 biological replicates) of three independent experiments (for one experiment, 8 plants were scored for each genotype). Asterisks indicate significant differences by Student’s two-tailed *t*-test (∗*P* < 0.05, ∗∗*P* < 0.01, ∗∗∗*P* < 0.001, ∗∗∗∗*P* < 0.0001). Different letters indicate statistically significant differences by two-way ANOVA and Tukey’s HSD (*P* < 0.05).

### Loss of *RAP2.6* gene is associated with metabolic remodeling during submergence

To determine whether RAP2.6-dependent submergence sensitivity was associated with altered metabolism, we performed untargeted metabolite profiling of four-week-old Col-0, *35S::RAP2.6*, and *rap2.6-1* mutant plants maintained in air or subjected to 24 h of dark submergence. Principal component analysis separated samples by treatment and genotype in both positive- and negative-ionization datasets (Fig. 2A, B; Figs. S3 and S4). PC1 and PC2 explained 63.1% and 11.6% of the variance, respectively, in positive mode and 72.9% and 7.9% in negative mode. Supervised PLS-DA and hierarchical clustering likewise indicated genotype- and treatment-dependent differences among detected features (Fig. 2C, D; Fig. S4). In positive polarity, metabolites such as steroidal glycoside, monogalactosyldiacylglycerol (MGDG) O-26:3_2:0, garcinolic acid, 17α-hydroxyprogesterone, and cryptotanshinone contributed strongly to class separation, whereas in negative mode the model was dominated by lipid- and acyl-related features, including phosphatidic acid (PA) 3:0_24:0, sulfoquinovosyl-diacylglycerol (SQDG) 23:0_18:4, hexosylceramide (HexCer) 25:1;2O/19:1, PS 10:0_10:0, and abruquinone B (Fig. 2C, D). Some key membrane-repair metabolites and lipids showed notable changes in *rap2.6-1* plants, including lysophosphatidylcholine (LPC) O-20:0, PA 3:0_24:0, phosphatidylserine (PS) 10:0_10:0, phosphatidylglycerol (PG) 6:0_24:4, MGDG O-26:3_2:0, and steroidal glycoside (Fig. S4A, B). In contrast, Col-0 and *35S::RAP2.6* plants exhibited membrane-damage lipids, including dilysocardiolipin (DLCL) 16:2_18:3, hemibismonoacylglycerophosphate (HBMP) 16:0_16:4_14:0, HexCer 25:1;2O/19:1, and SQDG 23:0_18:4 (Fig. S4A, B). Moreover, the volcano plot was analyzed collectively to demonstrate the metabolic changes in the air and sub groups across both ionization modes (Fig. S4C–H). Overall, the observed upregulated metabolites are corosolic acid, cryptotanshinone, pyridaben, and coniferin (Fig. S4C–H). Among these, corosolic acid is regarded as a significant, robust, and treatment-responsive metabolite, while lovastatin, LPC O-20:0, diacylglycerol (DG) 17:0_19:0, and triacylglycerol (TG) 8:0_8:0_28:3 are considered downregulated metabolites. *RAP2.6* overexpression showed the strongest positive-mode induction, especially cryptotanshinone, pyridaben, coniferin, and bonaniol A, while these metabolites redistributed in *rap2.6-1* knockout (Fig. S4C–H). In negative polarity, *rap2.6-1* plants upregulated metabolites such as eserine and N-acyl glycyl serine (NAGlySer) 18:3/24:2, compared with *35S::RAP2.6* (Fig. S4C–H). Collectively, several lipid-related features differed strongly between genotypes after submergence. Features annotated as PA, PS, PG, MGDG, LPC, N-acyl amino acids, and other lipid species were enriched in *rap2.6-1* relative to *35S::RAP2.6*, whereas the overexpression line accumulated a distinct set of lipid-associated features (Fig. S4).

**Fig. 2.**
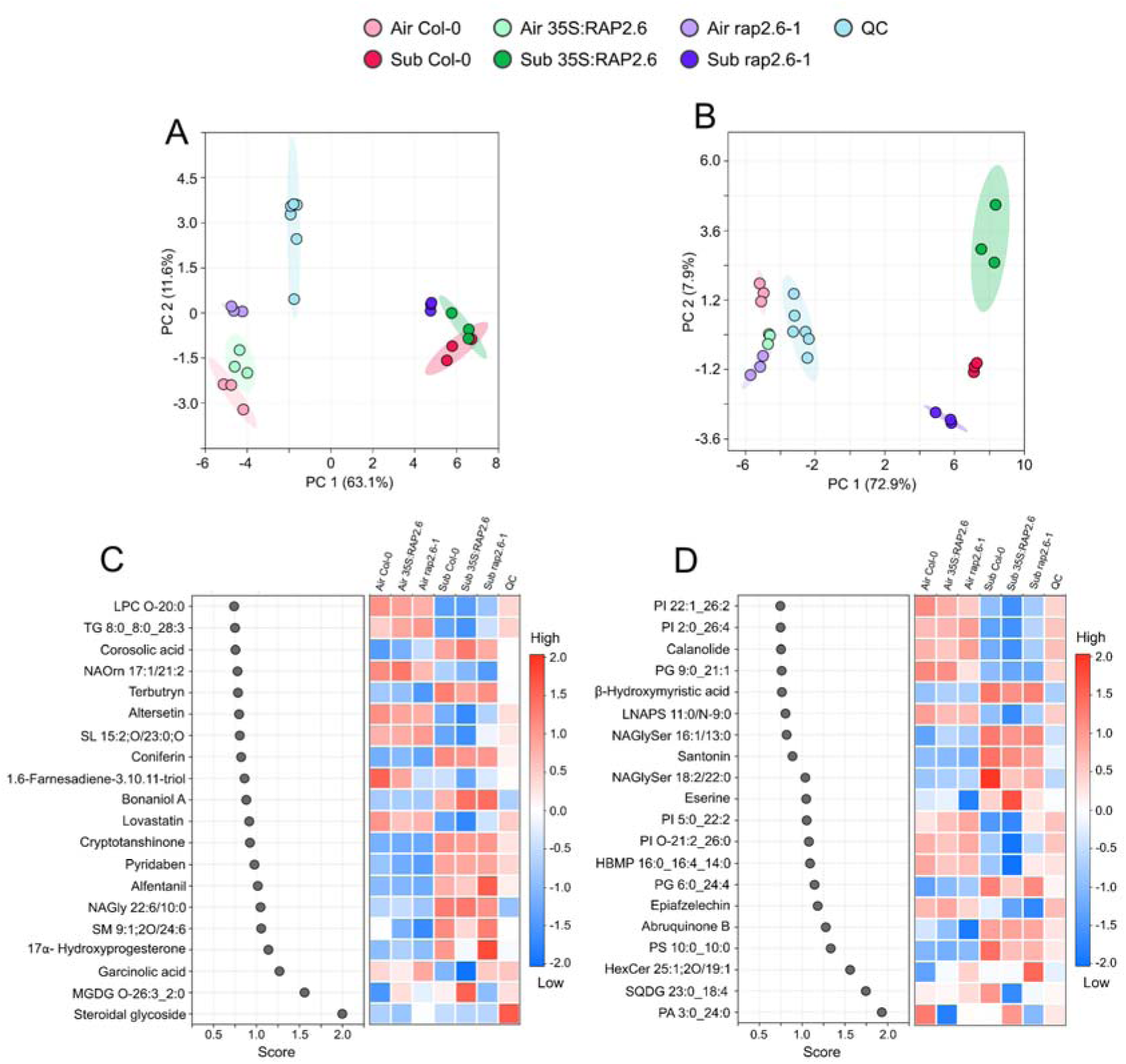
Metabolic profiling of RAP2.6 under submergence stress. Principal Component Analysis (PCA) of metabolites in Col-0, *35S::RAP2.6*, and *rap2.6-1* plants after 1 day of dark submergence or air, in (A) [M+H]^+^ and (B) [M-H]^-^ ion modes. Partial Least Squares – Discriminant Analysis (PLS-DA) of the top-ranked 20 metabolites based on variable importance in projection (VIP) score indicates metabolite discriminations among sample groups for (a) [M+H]^+^ and (b) [M-H]^-^ ion modes. Dots indicating higher scores correspond to greater contributions to metabolite identification. The colored map on the right shows the relative abundances of each metabolite across the control and treatment samples.

### RAP2.6 alters ethylene and nitric oxide homeostasis during submergence

Under control conditions, ethylene production and transcript abundance of the ethylene biosynthetic genes *1-AMINOCYCLOPROPANE-1-CARBOXYLATE OXIDASE 4* (*ACO4*) and *ACO5* were low and broadly similar among Col-0, *35S::RAP2.6*, and *rap2.6-1*. Following 24 h of dark submergence, ethylene production and expression of *ACO4* and *ACO5* significantly increased in Col-0 and *rap2.6-1* but remained lower in *35S::RAP2.6* (Fig. 3A-C). DAF-FM fluorescence decreased after submergence in all genotypes, but *35S::RAP2.6* retained a higher signal than Col-0 and *rap2.6-1* (Fig. 3D). Relative to Col-0, the overexpression line also showed greater NR activity, higher *NITRATE REDUCTASE 1* (*NIA1*) and *NIA2* transcript abundance, and notably increased SNO content (Fig. 3E-H) under dark submergence. Thus, *RAP2.6* overexpression is associated with relatively high NO/SNO-related signals and attenuated ethylene accumulation during submergence.

**Fig. 3.**
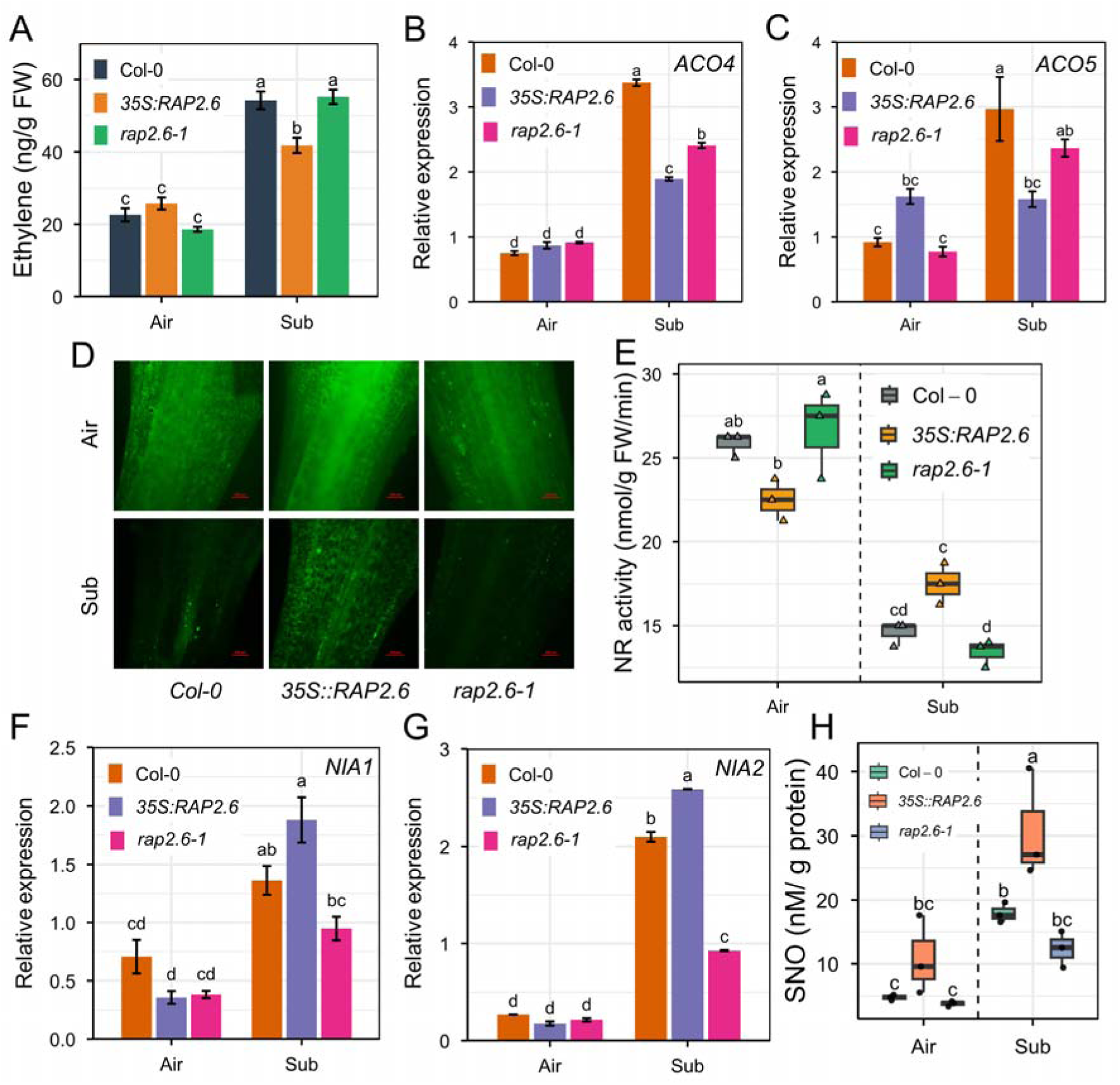
Analysis of nitric oxide and ethylene accumulation in RAP2.6 under submergence. (A) Ethylene levels in Col-0, *35S::RAP2.6*, and *rap2.6-1* plants exposed to air or dark submergence for 1 day. Bars represent means ± SE (*n* = 3) of three independent experiments (for one experiment, 4 plants were used for each genotype). (B, C) Relative gene expression of the ethylene biosynthesis gene, *ACO4* and *ACO5*, in Col-0, *35S::RAP2.6*, and *rap2.6-1* plants exposed to air or dark submergence for 24 hours (means ± SE, *n* = 3 biological replicates). (D) Representative fluorescence images visualizing NO using the fluorescent probe DAF-FM^TM^ diacetate, in Col-0, *35S::RAP2.6*, and *rap2.6-1* plant leaves exposed to air or dark submergence for 1 day. (E) Nitrate reductase (NR) activity, (F, G) relative gene expression of *NIA1* and *NIA2*, and (H) S-nitrosothiols (SNO) content in Col-0, *35S::RAP2.6*, and *rap2.6-1* plants exposed to air or dark submergence for 1 day (means ± SE, *n* = 3 biological replicates). Different letters indicate statistically significant differences by two-way ANOVA and Tukey’s HSD (*P* < 0.05).

### GSNOR1-dependent GSNO/SNO homeostasis determines RAP2.6-associated submergence tolerance

To distinguish the contribution of NO-related signaling from ethylene accumulation, we first crossed *RAP2.6* lines with *noa1*, which has reduced endogenous NO. Although *noa1* and the corresponding *RAP2.6 noa1* combinations accumulated more ethylene during submergence, they remained highly sensitive to prolonged dark submergence and exhibited increased electrolyte leakage (Figs. S5 and S6). Lower NO, accompanied by higher ethylene, was therefore insufficient to reproduce the tolerant phenotype of *rap2.6-1*.

We next examined GSNOR1 activity, which regulates GSNO/SNO homeostasis. The *gsnor1-1* line, characterized by increased GSNOR activity and a reduced SNO pool, showed greater survival under dark submergence and ethanol treatment, whereas *gsnor1-3*, which has reduced GSNOR activity and elevated SNO, was highly sensitive (Fig. S7). In double-mutant analyses, *gsnor1-1* substantially suppressed the submergence-sensitive phenotype of *35S::RAP2.6*, whereas *gsnor1-3* abolished the enhanced tolerance of *rap2.6-1* (Fig. 4A-D).

**Fig. 4.**
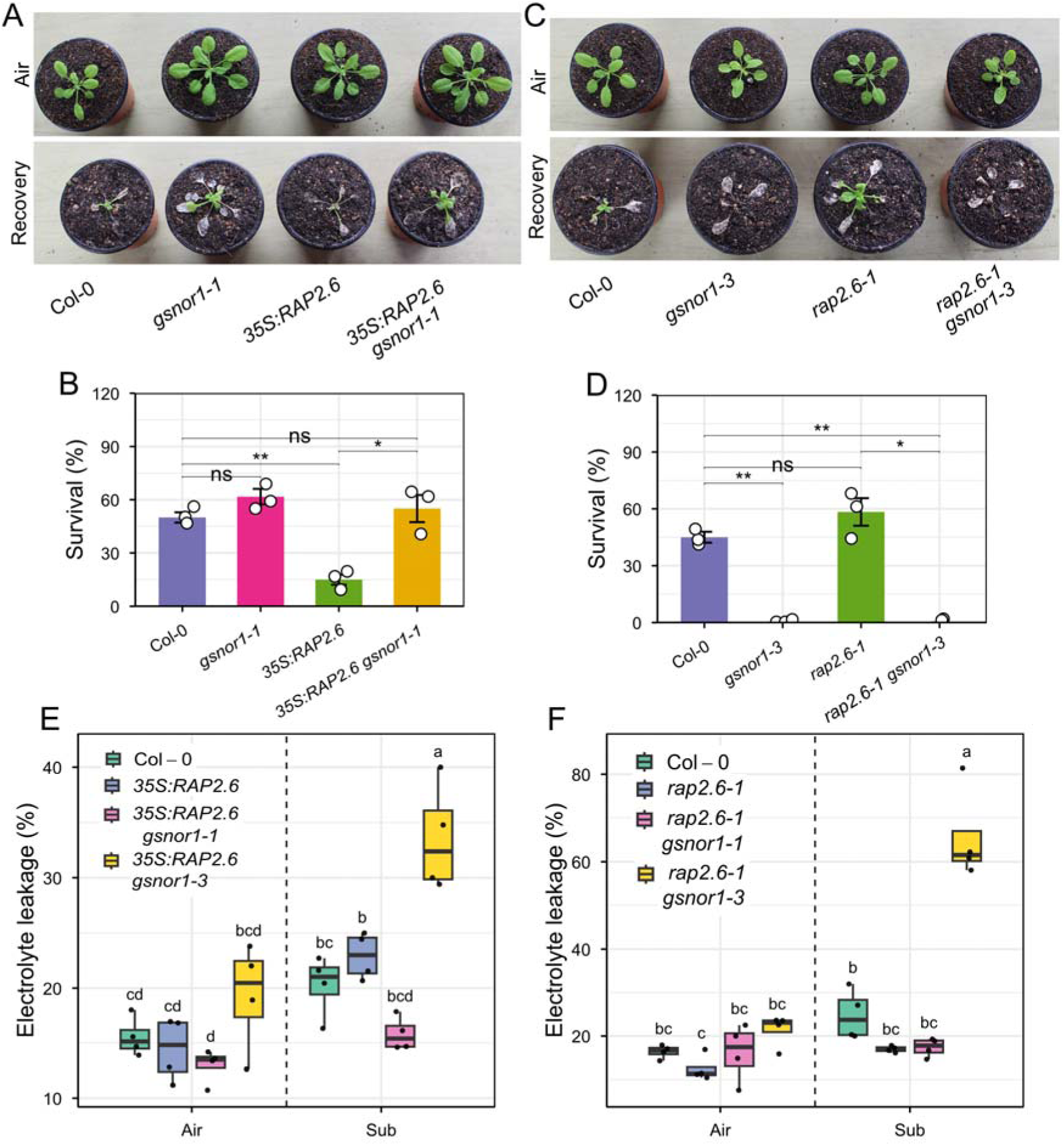
Evaluation of submergence tolerance in RAP2.6 expressing GSNOR1. (A, C) Phenotypes of Col-0, *gsnor1-1*, *35S::RAP2.6*, and *35S::RAP2.6 gsnor1-1*, and Col-0, *gsnor1-3*, *rap2.6-1*, and *rap2.6-1 gsnor1-3* plants were photographed before submergence (air) and after 5 days of dark submergence, followed by 7 days of recovery. (B, D) Survival rates were calculated after 7 days of recovery. Bars represent means ± SE (*n* = 3) of three independent experiments (for one experiment, >10 plants were scored for each genotype). (E, F) Electrolyte leakage in Col-0, *35S::RAP2.6*, *35S::RAP2.6 gsnor1-1*, and *35S::RAP2.6 gsnor1-3*, and Col-0, *rap2.6-1*, *rap2.6-1 gsnor1-1*, and *rap2.6-1 gsnor1-3* plants exposed to air or 2 days of dark submergence (means ± SE, *n* = 4 biological replicates for EL, *n* = 9 technical replicates from 3 biological replicates). Asterisks indicate significant differences by Student’s two-tailed *t*-test (∗*P* < 0.05, ∗∗*<u>P</u>* < 0.01, ∗∗∗*P* < 0.001, ∗∗∗∗*P* < 0.0001). Different letters indicate statistically significant differences by two-way ANOVA and Tukey’s HSD (*P* < 0.05).

Electrolyte leakage showed corresponding changes (Fig. 4E, F). These genetic interactions demonstrate that GSNOR1-dependent GSNO/SNO homeostasis is a major determinant of RAP2.6-associated submergence responses. Because the phenotypic relationships differ among measured outputs, we avoid assigning RAP2.6 and GSNOR1 to a single linear upstream-downstream hierarchy.

### GSNOR1 modifies RAP2.6-dependent NO/SNO and ethylene responses

*GSNOR1* genotype strongly modified RAP2.6-associated NO/SNO phenotypes. Under submergence, the low NO/SNO phenotype of *gsnor1-1* was retained in *35S::RAP2.6 gsnor1-1,* whereas the elevated SNO state associated with *gsnor1-3* was evident in the *rap2.6-1 gsnor1-3* background (Fig. 5A-D). Activity of NR was likewise influenced by *GSNOR1* genotype (Fig. S8). These observations indicate that GSNOR1-dependent GSNO/SNO homeostasis is required for the full effect of RAP2.6 on nitrosative signaling.

**Fig. 5.**
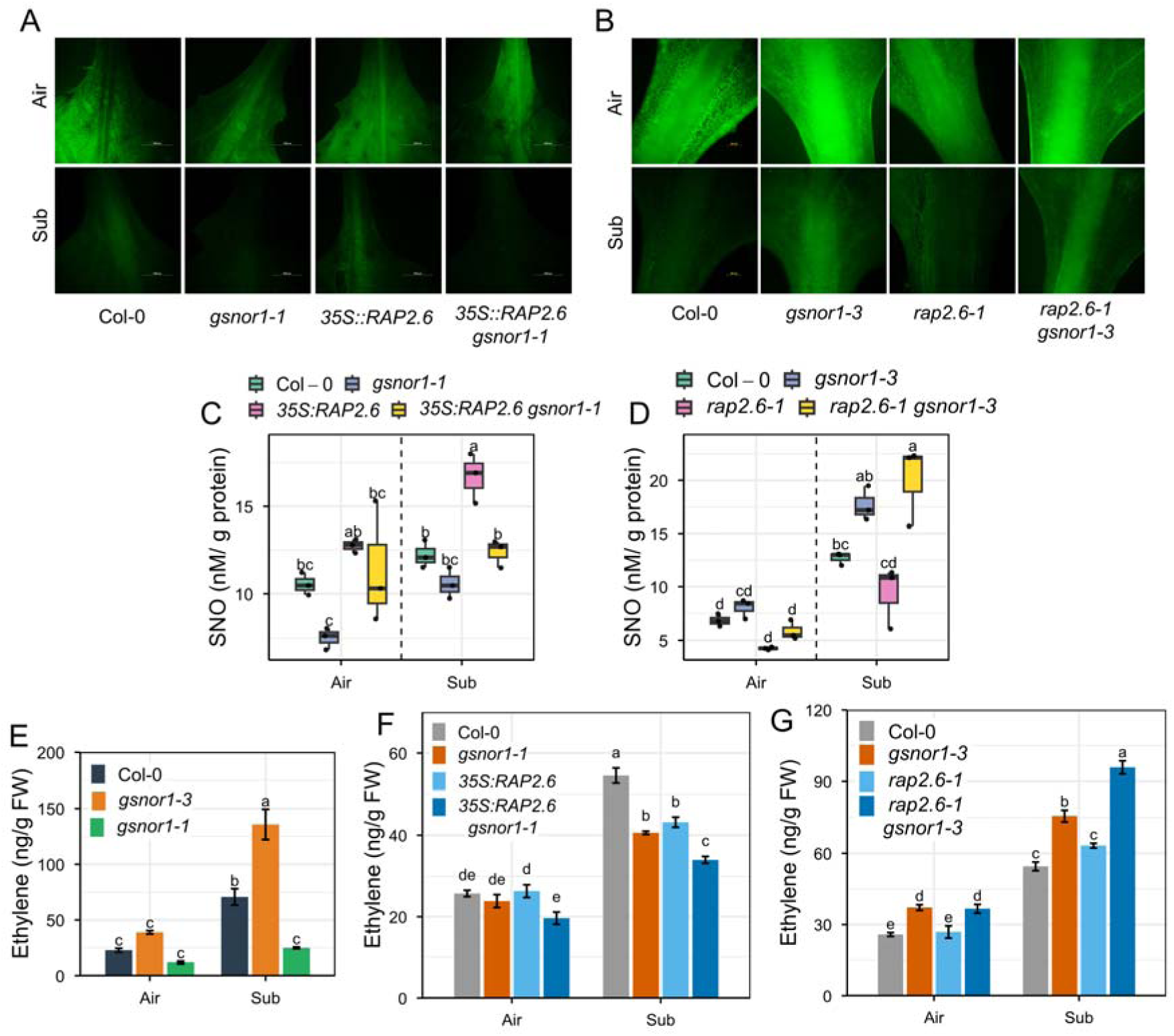
Analysis of nitric oxide and ethylene accumulation in RAP2.6 expressing GSNOR1 under submergence. (A, B) Representative fluorescence images visualizing NO using the fluorescent probe DAF-FM^TM^ diacetate, in Col-0, *gsnor1-1*, *35S::RAP2.6*, and *35S::RAP2.6 gsnor1-1*, *and* Col-0, *gsnor1-3*, *rap2.6-1*, and *rap2.6-1 gsnor1-3* plant leaves exposed to air or dark submergence for 1 day. (C, D) S-nitrosothiols (SNO) content in Col-0, *gsnor1-1*, *35S::RAP2.6*, and *35S::RAP2.6 gsnor1-1*, *and* Col-0, *gsnor1-3*, *rap2.6-1*, and *rap2.6-1 gsnor1-3* plants exposed to air or dark submergence for 1 day (means ± SE, *n* = 3 biological replicates). (E) Ethylene levels in Col-0, *gsnor1-3*, and *gsnor1-1* plants exposed to air or dark submergence for 1 day. (F) Ethylene levels in Col-0, *gsnor1-1*, *35S::RAP2.6*, and *35S::RAP2.6 gsnor1-1*, *and* (G) Col-0, *gsnor1-3*, *rap2.6-1*, and *rap2.6-1 gsnor1-3* plants exposed to air or dark submergence for 1 day. Bars represent means ± SE (*n* = 3) of three independent experiments (for one experiment, 4 plants were used for each genotype).Different letters indicate statistically significant differences by two-way ANOVA and Tukey’s HSD (*P* < 0.05).

The ethylene response was more complex. Despite elevated NO/SNO-associated signals, *gsnor1-3* accumulated more ethylene after submergence, and a similar increase was observed in the NO-overproducing *cue1-6* mutant (Fig. 5E; Fig. S9A). By contrast, *gsnor1-1* and *35S::RAP2.6 gsnor1-1* accumulated less ethylene (Fig. 5F), whereas both *35S::RAP2.6 gsnor1-3* and *rap2.6-1 gsnor1-3* showed high ethylene accumulation (Fig. 5G; Fig. S9B). Thus, the relationship between NO/SNO status and ethylene is context dependent, and the protective effect of GSNOR1 cannot be explained solely by a simple inverse relationship between ethylene and NO.

### RAP2.6 and GSNOR1 modulate transcriptional outputs associated with O_2_ sensing

Because NO and GSNO/SNO signaling have been linked to ERFVII turnover, we examined transcripts encoding components of the PCO N-degron pathway. Plant cysteine oxidase (PCO) 1 (*PCO1*) and *PCO2* transcript abundance increased in *35S::RAP2.6* and was reduced in *rap2.6-1* after 6 h of dark submergence (Fig. 6B, C). GSNOR1 genotype modified these RAP2.6-associated transcriptional responses (Fig. 6D-G). Transcripts of *RAP2.12*, *RAP2.3*, *HRE1*, *ALCOHOL DEHYDROGENASE 1* (*ADH1*), and *PYRUVATE DECARBOXYLASE 1* (*PDC1*) also differed among *RAP2.6* genotypes (Fig. S10). Because these measurements assess transcript abundance rather than PCO enzymatic activity or ERFVII protein stability, they indicate that RAP2.6 and GSNOR1 influence transcriptional outputs associated with the O_2_-sensing network but do not establish direct regulation of N-degron activity.

**Fig. 6.**
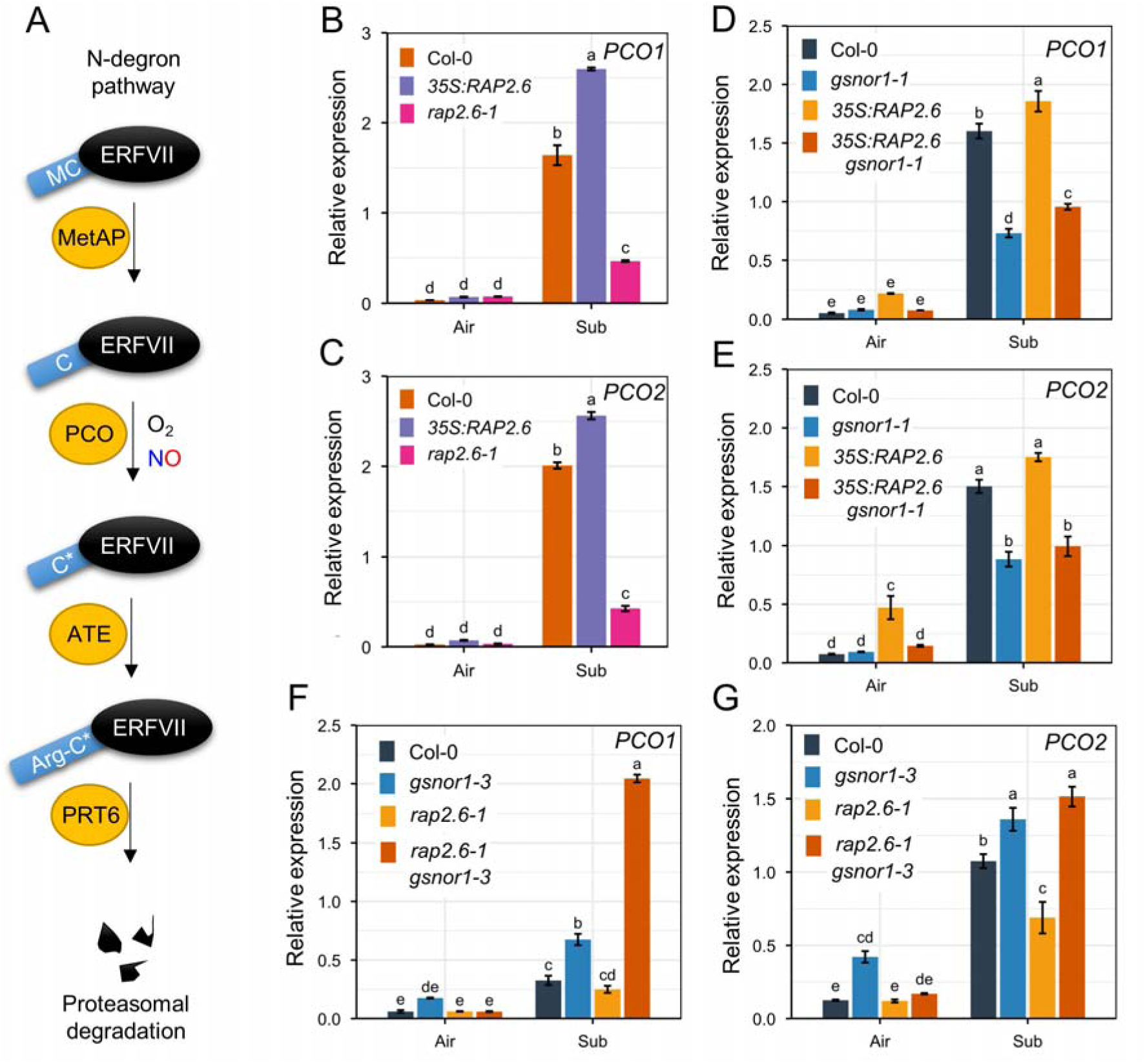
Relative gene expression analysis of enzymes involved in the N-degron under submergence. (A) Model mechanism of the ERFVII degradation via the N-degron pathway. (B, C) Relative gene expression related to the N-degron pathway, *PCO1* and *PCO2* in Col-0, *35S::RAP2.6*, and *rap2.6-1* plant leaves exposed to air or dark submergence for 24 hours (means ± SE, *n* = 3 biological replicates). Relative gene expression related to the N-degron pathway, *PCO1* and *PCO2*, in (D, C) Col-0, *gsnor1-1*, *35S::RAP2.6*, and *35S::RAP2.6 gsnor1-1*, *and* (F, G) Col-0, *gsnor1-3*, *rap2.6-1*, and *rap2.6-1 gsnor1-3* plant leaves exposed to air or dark submergence for 24 hours (means ± SE, *n* = 3 biological replicates). Different letters indicate statistically significant differences by two-way ANOVA and Tukey’s HSD (*P* < 0.05).

### GSNOR1 modifies RAP2.6-dependent ABA signaling during submergence

*RAP2.6* overexpression increased the transcript abundance of the ABA biosynthesis/signaling-associated genes *NINE-CIS-EPOXYCAROTENOID DIOXYGENASE 3* (*NCED3*), *SnRK2.6*, *CDK8*, *COR15A*, and *RD29A* after 6 h of submergence, whereas expression was generally lower or unchanged in *rap2.6-1* relative to Col-0 (Fig. 7A-E). GSNOR1 genotype strongly modified these responses: expression was reduced in *35S::RAP2.6 gsnor1-1* and enhanced in *gsnor1-3* and *rap2.6-1 gsnor1-3* (Fig. 7F, G). Consistent with this transcriptional response, *gsnor1-3* and its *RAP2.6* double-mutant combinations were hypersensitive to exogenous ABA (Fig. S11A-C). Endogenous ABA nevertheless declined after submergence across genotypes, with additional genotype-dependent differences (Fig. S11D, E). These results indicate that RAP2.6 and GSNOR1 primarily modify ABA responsiveness/signaling under submergence rather than simply increasing ABA abundance.

**Fig. 7.**
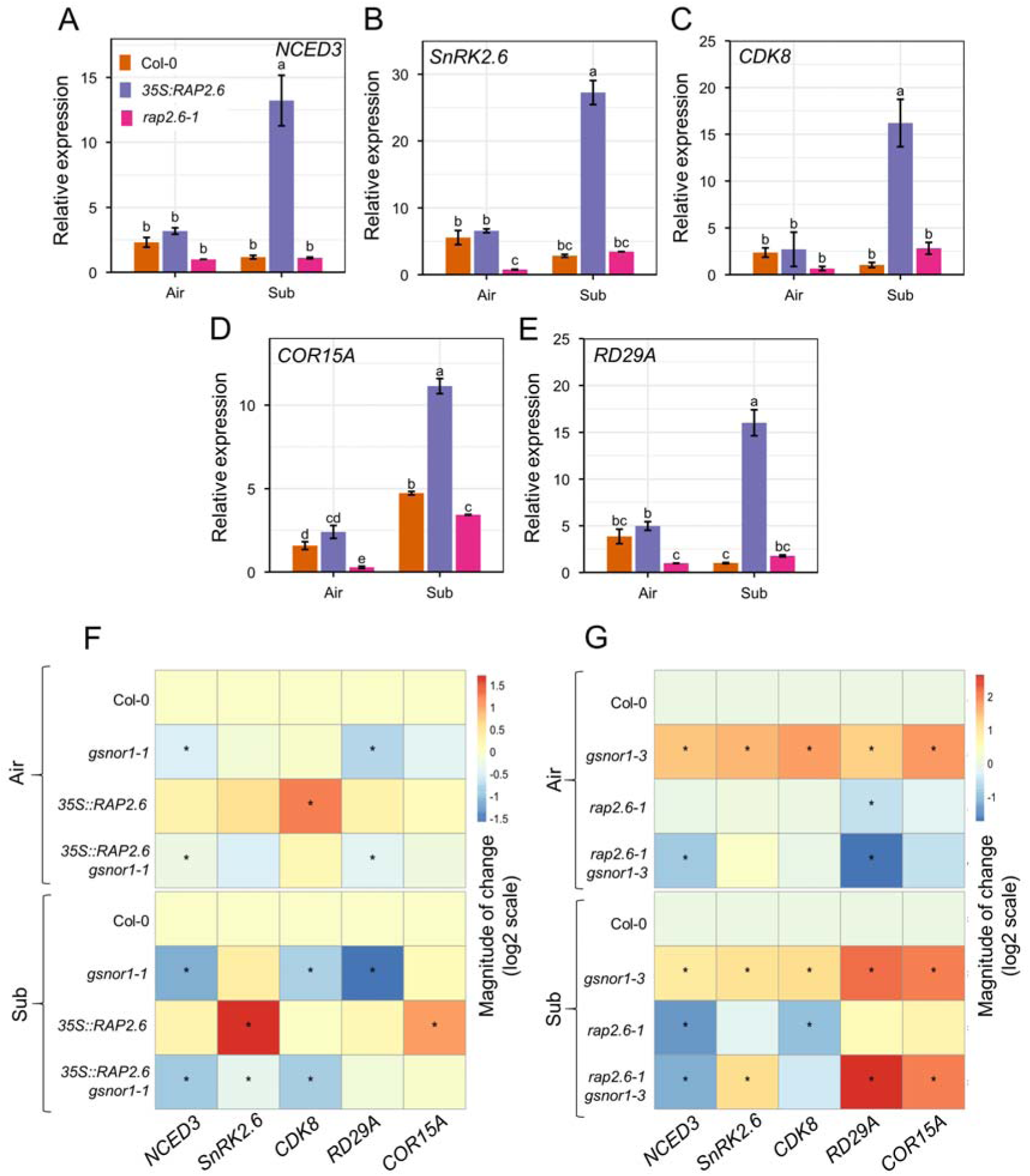
Relative gene expression analysis of ABA signaling under submergence. (A–E) Relative gene expression related to the ABA signaling, *NCED3*, *SnRK2.6*, *CDK8*, COR15A, and *RD29A*, in Col-0, *35S::RAP2.6*, and *rap2.6-1* plant leaves exposed to air or dark submergence for 24 hours (means ± SE, *n* = 3 biological replicates). Heatmap showing the relative gene expression related to the N-degron pathway, *NCED3*, *SnRK2.6*, *CDK8*, COR15A, and *RD29A*, in (F) Col-0, *gsnor1-1*, *35S::RAP2.6*, and *35S::RAP2.6 gsnor1-1*, *and* (G) Col-0, *gsnor1-3*, *rap2.6-1*, and *rap2.6-1 gsnor1-3* plant leaves exposed to air or dark submergence for 24 hours (means ± SE, *n* = 3 biological replicates). Different letters indicate statistically significant differences by two-way ANOVA and Tukey’s HSD (*P* < 0.05). Asterisks indicate significant differences by Student’s two-tailed *t*-test (∗*P* < 0.05, ∗∗*P* < 0.01, ∗∗∗*P* < 0.001, ∗∗∗∗*P* < 0.0001).

### GSNOR1 suppresses RAP2.6-associated oxidative damage and carbon depletion

Dark submergence increased H_2_O_2_ accumulation and malondialdehyde (MDA) content, with the strongest oxidative phenotype in *35S::RAP2.6* and lower levels in *rap2.6-1* (Fig. 8A-C). catalase (CAT), ascorbate peroxidase (APX), and peroxidase (POD) activities increased in *rap2.6-1* but were reduced in *RAP2.6*-overexpressing plants (Fig. 8D-F), consistent with enhanced antioxidant capacity in the loss-of-function mutant. In the genetic combinations, *gsnor1-1* suppressed H_2_O_2_ accumulation and oxidative damage in the *35S::RAP2.6* background, whereas *gsnor1-3* increased oxidative stress in the *rap2.6-1* background (Fig. 8G-L). GSNOR1 genotype also modified total soluble sugar and carbohydrate pools, where *gsnor1-1* restored these reserves in *35S::RAP2.6* but *gsnor1-3* reduced them in *rap2.6-1* (Fig. S12). Together, these data link GSNOR1-dependent redox homeostasis with antioxidant capacity and maintenance of carbon reserves during submergence.

**Fig. 8.**
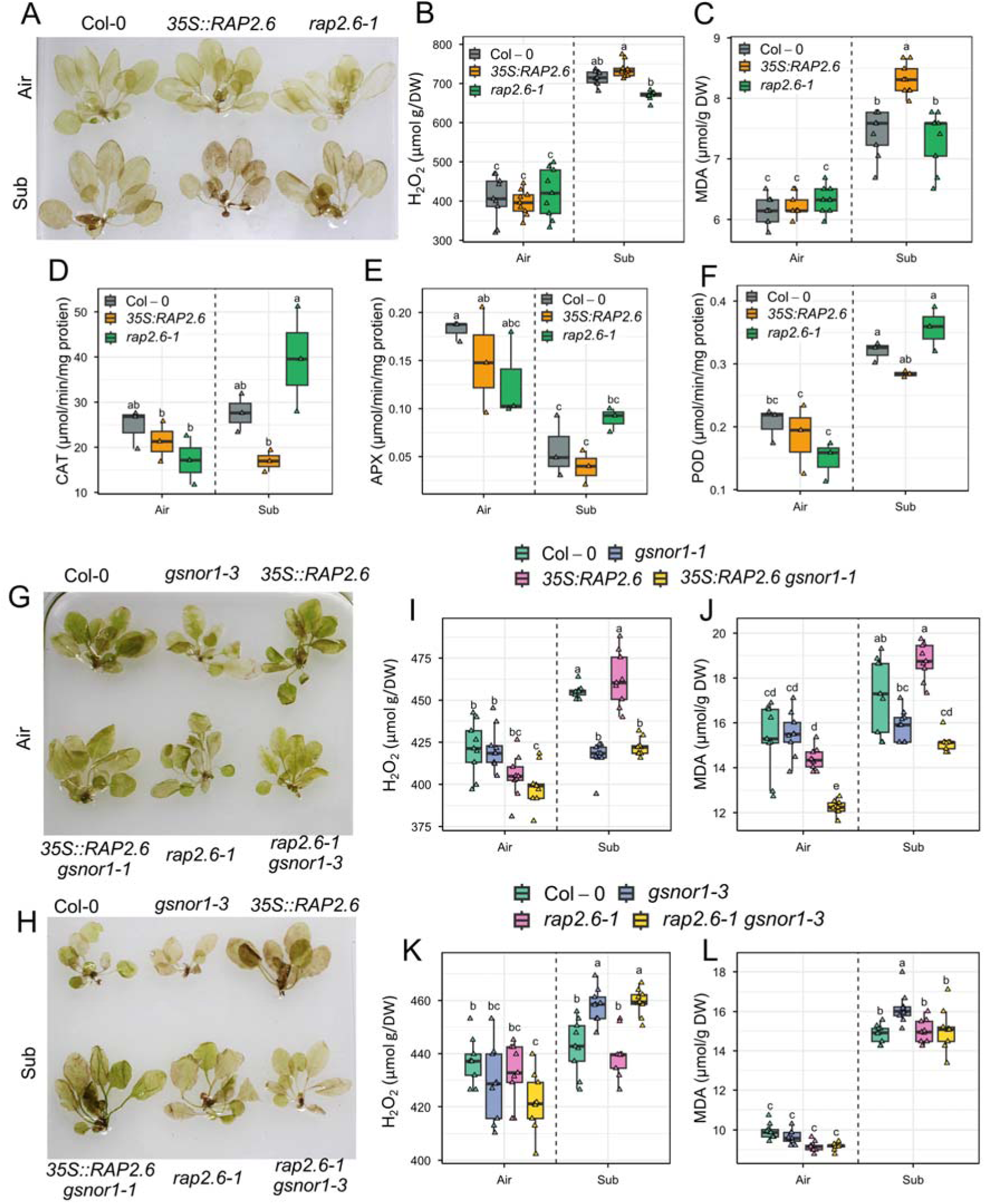
Analysis of oxidative damage and antioxidant activities under submergence. (A) DAB staining was used to assess the accumulation of H_2_O_2_ in the leaves of Col-0, *35S::RAP2.6*, and *rap2.6-1* plants exposed to air or dark submergence for 2 days. (B, C) Levels of H_2_O_2_ and MDA, in Col-0, *35S::RAP2.6*, and *rap2.6-1* plants exposed to air or dark submergence for 2 days (means ± SE, *n* = 9 technical replicates from 3 biological replicates). (D–F) Antioxidant activity of CAT, APX, and POD in Col-0, Col-0, *35S::RAP2.6*, and *rap2.6-1* plants exposed to air or dark submergence for 2 days (means ± SE, *n* = 3 biological replicates). (G, H) DAB staining was used to assess the accumulation of H_2_O_2_ in the leaves of Col-0, *gsnor1-3*, *35S::RAP2.6*, *35S::RAP2.6 gsnor1-1*, *rap2.6-1*, *rap2.6-1 gsnor1-3* plants exposed to air or dark submergence for 2 days. (I, J) Levels of H_2_O_2_ and MDA, in Col-0, *gsnor1-1*, *35S::RAP2.6*, *35S::RAP2.6 gsnor1-1*, plants exposed to air or dark submergence for 2 days (means ± SE, *n* = 9 technical replicates from 3 biological replicates). (K, L) Levels of H_2_O_2_ and MDA, in Col-0, *gsnor1-3*, *rap2.6-1*, *rap2.6-1 gsnor1-3*, plants exposed to air or dark submergence for 2 days (means ± SE, *n* = 9 technical replicates from 3 biological replicates). Different letters indicate statistically significant differences by two-way ANOVA and Tukey’s HSD (*P* < 0.05). DW: freeze-dry weight.

## Discussion

During flooding, plants favor the fermentation process over oxidative phosphorylation for ATP production. To adapt to and survive low O_2_ levels, plants also employ morphological acclimation together with signaling networks, particularly by stabilizing the ERFVII master TFs in the nucleus (Wang et al., 2025). However, the activation of PCO enzymes and the presence of NO favor the Arg-Cys N-terminal proteolytic degradation of ERFVII via the N-degron pathway (Giuntoli and Perata, 2017). Beyond ERFVII, the broader AP2/ERF family remains underexamined in O_2_-sensing mechanisms and its links to NO signaling. In this study, we investigated a group X ERF transcription factor RAP2.6, which is reported to be involved in multiple stress signaling processes but not in responses to low O_2_.

Our results identify RAP2.6 as a negative regulator of Arabidopsis tolerance to O_2_ limitation. *RAP2.6* overexpression reduced survival after dark submergence and increased sensitivity to hypoxia and ethanol, whereas loss of *RAP2.6* produced the opposite phenotype (Fig. 1). This pattern extends the contribution of AP2/ERF transcription factors to flooding responses beyond the canonical ERFVII O_2_ sensors. The phenotype is also consistent with recent evidence that other non-ERFVII family members can modulate hypoxia tolerance (Liao et al., 2026). Importantly, however, RAP2.6 lacks the characteristic N-terminal cysteine that confers direct O_2_-dependent stability on ERFVIIs. RAP2.6 should therefore be viewed as a regulator of low-O_2_ acclimation rather than as a canonical O_2_ sensor.

The physiological phenotype of *rap2.6-1* was accompanied by maintenance of photosynthetic pigments and soluble sugars, lower electrolyte leakage, and extensive metabolite remodeling. Lipid-associated features assigned to PA, PS, PG, MGDG, LPC, and N-acyl amino acids differed strongly between *RAP2.6* overexpressing and knockout plants after submergence (Fig. 2). These lipid classes have established roles in membrane organization, signaling, and stress acclimation (Sakurai et al., 2003; Bogos et al., 2010; Yamaoka et al., 2021; Yao et al., 2024). Nevertheless, untargeted metabolite annotations vary in confidence, and several compound assignments in our dataset require orthogonal validation. We therefore interpret the current dataset as evidence for RAP2.6-dependent metabolic and lipid remodeling rather than direct proof that individual annotated metabolites mediate membrane repair or injury.

A major finding is the strong genetic interaction between RAP2.6 and GSNOR1-dependent GSNO/SNO homeostasis. Increasing GSNOR activity in *gsnor1-1* suppressed the submergence sensitivity of *RAP2.6*-overexpressing plants, whereas loss of GSNOR1 activity in *gsnor1-3* abolished the enhanced tolerance of *rap2.6-1* (Fig. 4). Under hypoxic conditions, *gsnor1-3* plants show similar hypersensitivity (Zhan et al., 2018). In our study, we found that these reciprocal genetic effects were accompanied by changes in NO/SNO-associated signals, oxidative stress, ABA responsiveness, and carbon reserves. Because GSNOR1 primarily regulates GSNO/SNO homeostasis rather than serving simply as a determinant of instantaneous free-NO concentration, the contrasting phenotypes of *noa1* and *gsnor* mutants are biologically informative. Lower NO in *noa1* was associated with increased ethylene but did not improve survival, whereas increased GSNOR activity conferred tolerance despite a different ethylene profile (Fig. 4; Fig. S6). Thus, NO abundance, GSNO/SNO homeostasis, and ethylene accumulation cannot be treated as interchangeable readouts of a single linear pathway.

The ethylene data further emphasize this context dependence. Early ethylene accumulation can promote NO depletion through phytoglobin-dependent mechanisms and enhance hypoxia acclimation (Hartman et al., 2019; van der Wijk et al., 2026). Consistent with this framework, *35S::RAP2.6* plants accumulated less ethylene and retained higher NO/SNO-associated signals than Col-0 during submergence. However, *gsnor1-3* accumulated both high SNO-associated signals and high ethylene, and *cue1-6* showed a similar increase in ethylene (Fig. 5). NO-dependent *S*-nitrosation can also modify enzymes involved in ethylene biosynthesis under other stresses (Liu et al., 2023). Collectively, these observations suggest that the effect of NO-related signaling on ethylene depends on the biochemical form and cellular context of the NO pool, including GSNO/SNO turnover. Direct identification of *S*-nitrosated targets in the ethylene pathway will be required to resolve this mechanism.

RAP2.6 and GSNOR1 also altered transcriptional outputs associated with the PCO-ERFVII pathway. *PCO1* and *PCO2* transcripts were higher in *35S::RAP2.6* and lower in *rap2.6-1*, and these responses were modified by GSNOR1 genotype (Fig. 6). Hypoxia-responsive transcripts, including *ADH1* and *PDC1*, were paradoxically elevated in the submergence-sensitive *RAP2.6*-overexpressing line. This dissociation between marker-gene induction and survival is not unprecedented and reinforces the distinction between transcriptional activation of hypoxia-responsive genes and successful acclimation (Hess et al., 2011; Zhan et al., 2018; Liu et al., 2022). Because the present experiments did not measure PCO enzymatic activity or ERFVII protein stability, we cannot conclude that RAP2.6 directly controls N-degron-mediated O_2_ sensing. Instead, our data indicate that RAP2.6 and GSNOR1 influence transcriptional outputs connected to this network. Protein-level analyses of ERFVIIs and direct measurements of PCO activity will be needed to establish whether the N-degron pathway is mechanistically altered.

ABA signaling provides an additional point of convergence. Studies reveal that RAP2.6 modulates ABA responses in *A. thaliana* under drought and salt stress (Zhu et al., 2020; Xuan et al., 2026). In our study, *RAP2.6* overexpression enhanced the expression of *NCED3*, *SnRK2.6*, *CDK8*, *COR15A*, and *RD29A* during submergence, and GSNOR1 genotype reversed both these transcriptional responses and ABA sensitivity (Fig. 7). Endogenous ABA nevertheless declined during submergence. The data therefore support altered ABA responsiveness rather than a simple RAP2.6-dependent increase in ABA accumulation.

Enhanced ABA signaling could favor growth restraint and metabolic conservation, but the present experiments do not directly demonstrate ABA-induced metabolic arrest. We therefore interpret ABA as one component of the RAP2.6-GSNOR1 response network whose contribution to survival will require targeted genetic or pharmacological dissection.

Finally, the contrasting oxidative and carbon phenotypes provide a physiological explanation for the survival differences. *RAP2.6* overexpression was associated with greater H_2_O_2_ and MDA accumulation, reduced antioxidant enzyme activities, and depletion of soluble sugars and carbohydrates, whereas *rap2.6-1* maintained stronger antioxidant capacity and carbon reserves (Fig. 8; Fig. S12). Increased GSNOR activity reversed several of these defects in the *35S::RAP2.6* background. Zhang et al. (2026) show that tobacco (*Nicotiana tabacum*) *GSNOR* overexpression lines enhance enzymatic and non-enzymatic activities, thereby improving ROS homeostasis and salt tolerance. We therefore propose that GSNOR1-dependent GSNO/SNO homeostasis buffers RAP2.6-associated nitro-oxidative imbalance and helps preserve the metabolic resources required for recovery from prolonged submergence.

Together, the data support a model in which RAP2.6 functions as a non-ERFVII negative regulator of submergence tolerance whose effects are strongly conditioned by GSNOR1-dependent redox homeostasis (Fig. 9). Rather than placing RAP2.6, GSNOR1, NO, ethylene, and ABA in a single linear cascade, we propose that their interactions generate multiple, partially separable outputs involving nitrosative signaling, hormone responsiveness, oxidative stress, and carbon metabolism. This framework reconciles the apparently divergent NO/SNO and ethylene phenotypes and provides testable hypotheses for future studies of protein S-nitrosation, ERFVII stability, and metabolic acclimation under O_2_ limitation.

**Fig. 9.**
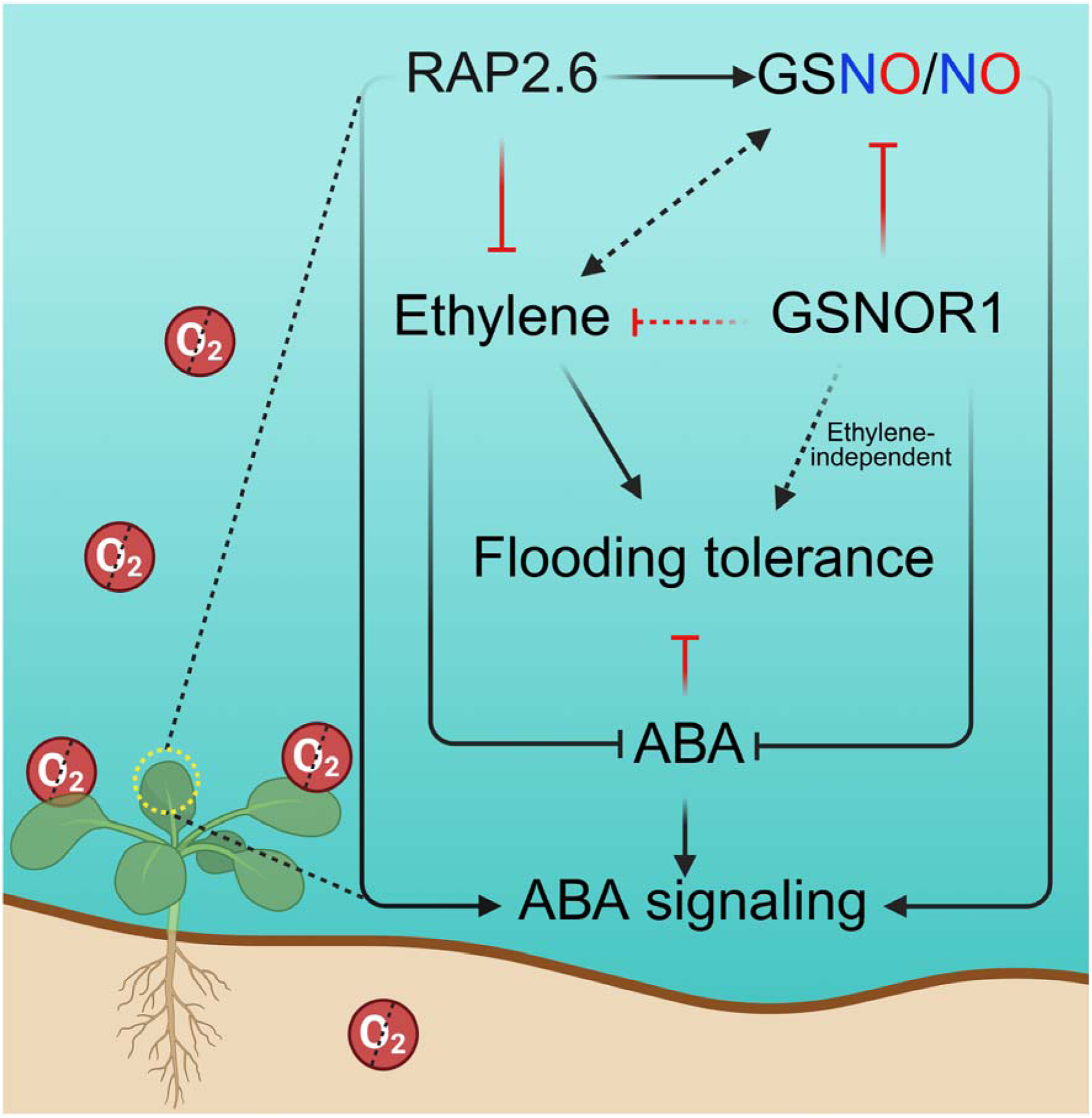
A putative working model of RAP2.6-mediated submergence sensitivity and its associated signaling with NO, ethylene, and ABA. RAP2.6 positively regulates endogenous NO levels in *Arabidopsis thaliana*, while limiting the ethylene accumulation and increasing ABA responses, thereby impairing the flooding tolerance. GSNOR1-driven redox balance in RAP2.6 altered the low O_2_ sensitivity, but the relationship between NO/SNO status and ethylene is context dependent, and the GSNOR1-mediated tolerance cannot be explained solely by a simple inverse relationship between ethylene and NO. NO, nitric oxide; GSNO, *S*-nitrosogluthathione; GSNOR1, *S*-nitrosogluthathione reductase 1; ABA, abscisic acid; O_2_, oxygen. Arrow-headed and blunt-ended (T-shaped) lines indicate positive and negative regulation, respectively. The dotted arrow indicates a possible direct regulation.

## Conclusions

Our findings identify RAP2.6 as a negative regulator of submergence tolerance in Arabidopsis. Loss of *RAP2.6* improves survival under O_2_-limiting conditions and is associated with reduced membrane injury, maintenance of carbon reserves, enhanced antioxidant capacity, and broad metabolic remodeling. Conversely, *RAP2.6* overexpression alters NO/GSNO and ethylene homeostasis, enhances ABA-responsive signaling and oxidative damage, and compromises survival during submergence. Genetic analyses further reveal that GSNOR1-dependent GSNO/SNO homeostasis is a major determinant of RAP2.6-associated stress responses. Increased GSNOR1 activity suppresses the submergence sensitivity of *RAP2.6*-overexpressing plants, whereas loss of GSNOR1 abolishes the enhanced tolerance of *rap2.6-1* mutant plants. The effects of GSNOR1 cannot be explained solely by ethylene accumulation, indicating that redox regulation and hormonal signaling converge through partially distinct pathways. These results expand the low-O_2_ regulatory network beyond canonical ERFVIIs and establish a functional connection between RAP2.6 and GSNOR1-dependent redox signaling during submergence. Given its previously characterized roles in abiotic and biotic stress responses, RAP2.6 may represent a promising molecular target for engineering low-O_2_ resilience in crop species. Future studies will be valuable for determining which key mechanisms underlie the discrepancy between GSNO and NO in controlling ethylene signaling under limited O_2_ conditions.

## Materials and methods

### Plant materials and growth conditions

All mutants used in this study were in the *A. thaliana* Columbia (Col-0) background. Seeds of the T-DNA insertion line *rap2.6-1* (N301757) and the *35S::RAP2.6* overexpression line were kindly provided by Dr. Holger Bohlmann (University of Natural Resources and Life Sciences, Vienna, Austria) and were described previously (Ali et al., 2013). The GSNOR1 gain- and loss-of-function lines *gsnor1-1* and *gsnor1-3*, respectively, were described by Feechan et al. (2005). *cue1-6* (CS3168) and *noa1* (CS6511) used in this study were described previously (Guo et al., 2003; He et al., 2004). Double homozygous lines were generated by crossing *35S::RAP2.6* or *rap2.6-1* with *gsno1*-*1*, *gsnor1-3*, or *noa1*. After validation of heterozygous F1 plants, selfed F2 progeny were genotyped by PCR, and F3/F4 seeds from confirmed double homozygous lines were used in the experiments (Das et al., 2026). Seeds were germinated in horticultural soil (Bio Top Soil No. 1, Heungnong Trading Company, South Korea) at 22 ± 1°C under a 16-h-light/8-h-dark photoperiod. Two-week-old seedlings were transplanted into 7-cm-high × 7-cm-diameter pots and maintained under the same growth conditions.

### Submergence, hypoxia, and ethanol treatments

Submergence tolerance assays were performed as described by Licausi et al. (2011). Four-week-old plants were submerged in 23-cm-high plastic boxes and maintained in darkness, with rosettes positioned approximately 14 cm below the water surface. After 5 d of submergence, plants were returned to normal growth conditions for 7 d. Survival was scored based on the production of new leaves and continued growth during recovery. Three independent experiments were performed, with more than 10 plants per genotype in each experiment. Samples were collected at the indicated times. For hypoxia treatment, sterilized seeds were grown on half-strength Murashige and Skoog (MS) solid medium for 7 d and then placed for 24 h in a sealed jar containing an Oxoid AnaeroGen 2.5-L sachet (Thermo Scientific; OXAN0025A), following Sherwood et al. (2023). Green cotyledons were scored after 3 d of recovery. Ethanol assays were performed according to Chen et al. (2015) by growing sterilized seeds on half-strength MS medium with or without 100 mM ethanol under a 16-h-light/8-h-dark photoperiod.

### Measurement of photosynthetic pigments

Total chlorophyll, carotenoids, and pheophytin were measured in 80% acetone extracts by recording absorbance at 663, 645, and 470 nm using a Multiskan GO microplate spectrophotometer (Thermo Scientific). Pigment concentrations were calculated according to Arnon (1949), Lichtenthaler and Wellburn (1983), and Lichtenthaler (1987), with normalization to sample mass and extraction volume.

### Quantification of total soluble sugars and total carbohydrates

Total soluble sugar (TSS) and total carbohydrate contents were quantified according to Somogyi (1952) and DuBois et al. (1956), respectively. Freeze-dried samples were extracted in 80% ethanol and centrifuged at 14,000 rpm for 20 min at 4°C. For TSS determination, the supernatant was diluted 1:4 with distilled water and mixed with 2.5 mL anthrone reagent (5 mg anthrone per 2.5 mL 98% H_2_SO_4_), followed by incubation for 20 min at 90°C. Absorbance was measured at 620 nm after cooling. For total carbohydrate determination, the supernatant was diluted 1:9, mixed with 500 µL 5% phenol and 2.5 mL 98% H_2_SO_4_, incubated for 20 min at 90°C, and measured at 490 nm after cooling. Concentrations were calculated from glucose standard curves and expressed as mg g^−1^ dry weight.

### Metabolites analysis by UPLC-HRMS

The freeze-dried samples were homogenized using an OMNI Bead Ruptor Elite bead mill homogenizer (BioD Co., Ltd., Korea). The homogenization was conducted at 4 m/s for 4 cycles, where each consisting of 15 sec run followed by 10 sec dwell time using 4 beads in each tube. The homogenized sample (150 mg) was extracted with 1 mL of an ice-cold solvent mixture of ACN:MEOH:IPA:H2O (3:3:2:2). The mixture was sonicated on ice for 15 min to perform ultrasound-assisted extraction. Subsequently, the samples were vortexed using a thermal shaker (Allsheng, China) at 1,500 rpm for 90 min at 4°C, followed by centrifugation at 15,000 rpm for 10 min at 4°C. The resulting supernatants were transferred into LC vials for UPLC–MS/MS analysis. To assess analytical reproducibility, each sample was analyzed in three consecutive runs. In addition, a pooled quality control (QC) sample was prepared by combining 10 μL aliquots from each sample extract prior to LC–MS analysis. Five QC injections were distributed in total across the analytical batch to evaluate the consistency, stability, and reliability of the analytical method.

The untargeted metabolite identification was performed using the cyclic ion mobility quadrupole time-of-flight mass spectrometer (cIM–TOF–MS) coupled to an ACQUITY I-Class UPLC system (Waters Corporation, Milford, MA, USA). The chromatographic separation was conducted using ACQUITY UPLC CSH Phenyl-Hexyl column (2.1 mm × 100 mm, 1.7 μm particle size; Waters Corporation, Milford, MA) at column temperature of 40 °C. A flow rate of 0.25 mL/min was used during LCMS acquisition. The mobiles phases, water with 0.1% formic acid (A) and acetonitrile with 0.1% formic acid (B) were used. The gradient elution program was set as follows: 0 min, 1% B; 3 min, 1% B; 6 min, 20% B; 10 min, 40% B; 15 min, 70% B; 20 min, 75% B; 22 min, 80% B; 24 min, 85% B; 25 min, 90% B; 27 min, 1% B; 30 min, 1% B. The flow rate was maintained at 0.250 mL/min throughout the run.

Mass spectrometric analysis was conducted using a quadrupole cyclic ion mobility mass spectrometer (Waters Corporation, Wilmslow, UK) equipped with an electrospray ionization source operated in MSE continuum mode. The MS^E^ data acquisition were performed at both positive and negative mode by applying the low and high transfer collision energy of 4V and ramping 20∼50 V, respectively. The capillary voltage at 3.2 kV, cone voltage at 40 V, source temperature at 150°C, desolvation temperature at 600°C, desolvation gas flow at 600 L/h, source offset at 10 V and IM cell pressure at 2.26 mbar N_2_ were set. Nitrogen (N_2_) and helium (He) gases were used as cone and collision gases, respectively. The data was acquired within the range of m/z 50 – 1200 Da. Leucine Enkephalin was used as the lockspray reference compound for continuous mass calibration throughout the analysis, with reference ions at m/z 556.28 ([M+H]) and m/z 554.26 ([M–H]). The lockspray solution was infused at a concentration of 50 pg/mL and a flow rate of 10 μL/min.

### Data processing and establishment of in-house database

The raw LC–MS data was processed using Progenesis QI 3.0 software (Waters, Milford, MA, USA) by following the processing steps – normalization, peak alignment, experiment design setup, peak picking, deconvolution, and metabolite identification. The retention time was set to 0.5 − 28.0 min for peak picking section. For positive ionization mode, the following adduct ions were selected for deconvolution: [M+H]^+^, [M+NH_4_]^+^, [M+Na]^+^, [M+K]^+^, [M+Cl]^+^, [M+H−H_2_O]^+^, [M+ACN+H]^+^, [M+ACN+Na]^+^, [M+H+Na]^+^ and [2M+H]+. For negative ionization mode, the selected adducts included [M–H]^-^, [M–NH_4_]^−^, [M–Na]^−^, [M–K]^−^, [M– Cl]^−^, [M−H_2_O–H]^−^, [M+Na–2H]^−^, [M+K–2H]^−^ and [M–2H]^−^ ion adducts were selected for negative mode. Compound features were further characterized using the Progenesis MetaScope module by matching MS/MS spectra against MSP spectral libraries constructed from multiple databases, including the Vaniya/Fiehn Natural Products Library, HCD Natural Product Library, ReSpect Consortium Library, LipidBlast 2022, MetaboBASE, GNPS NIH Natural Products Library, PFP Natural Products Library, MassBank, and the In-house Antarctic Moss Library. For metabolite identification, precursor ion and fragment ion mass tolerances were both set to 10 ppm. Detected compounds were annotated based on composite identification criteria, including overall score, isotope similarity, fragmentation score, and mass error. In addition, the in-house Antarctic moss metabolite library was established and applied for both positive ([M+H]□) and negative ([M−H]□) ionization modes.

### Multivariate analysis

The raw LC–MS data were exported by containing the information of detected peaks – m/z value and peak intensities. The Metaboanalyst (v6.0, https://www.metaboanalyst.ca/) was used to further analyze the post-processing data. The data post-processing involved normalization by sum, log transformation, and Pareto scaling for multivariate statistical analysis.

### Ethylene measurement

Ethylene production was quantified after 1 d of dark submergence. Four rosettes from submerged or air-grown plants were gently blotted, weighed, and placed in 20-mL airtight vials. Vials were sealed and incubated in darkness for 2 h under the same temperature conditions used for plant growth. A 1-mL headspace sample was analyzed using an Agilent 6890N gas chromatograph equipped with a flame-ionization detector. Injector and detector temperatures were 200°C and 250°C, respectively, and the oven temperature was 80°C. Ethylene concentrations were calculated from a standard curve, normalized to fresh weight, and expressed as ng g^−1^ FW.

### Fluorescence detection of nitric oxide using DAF-FM

Leaf NO-associated fluorescence was assessed using DAF-FM diacetate (Thermo Fisher Scientific). Leaves were preincubated for 15 min in MES-KCl buffer (10 mM MES, 50 mM KCl, pH 6.15, 0.05% Tween-20) in darkness and then incubated with freshly prepared 10 µM DAF-FM diacetate for 40 min with gentle shaking. Samples were vacuum infiltrated for 15 min in darkness, washed four to five times with buffer, mounted on glass slides, and imaged using a Nikon ECLIPSE Ti2-U fluorescence microscope (excitation, 480 nm; emission, 500-540 nm). Samples without DAF-FM were used to estimate autofluorescence. Five biological replicates were analyzed using identical microscope settings. Because DAF-FM fluorescence is an indirect probe of NO-related reactive nitrogen species, results are referred to as NO-associated fluorescence rather than an absolute measurement of free NO concentration.

### Nitrate reductase activity

Nitrate reductase (NR) activity was quantified according to Samant et al. (2025) with minor modifications. Fresh tissue (200 mg) was homogenized in 400 µL extraction buffer containing 100 mM HEPES (pH 7.6), 15 mM MgCl_2_, 3.5 mM β-mercaptoethanol, 0.5% PVP, and 0.3% Triton X-100. After addition of 50 µL 0.5 M zinc acetate, samples were incubated on ice for 30 min and centrifuged at 14,000 rpm for 20 min at 4°C. Twenty microliters of supernatant was mixed with 80 µL assay buffer containing 100 mM HEPES-KOH (pH 7.6), 0.5 mM DTT, 50 mM MgCl_2_, 5 mM KNO_3_, and 0.2 mM NADH. Griess reagent (100 µL; Sigma-Aldrich) was added, samples were incubated for 10 min at room temperature, and absorbance was measured at 546 nm. NR activity was calculated using a NaNO_2_ standard curve and expressed as nmol nitrite g^−1^ FW min^−1^.

### RNA isolation and RT-qPCR analysis

Total RNA was extracted from 200 mg of liquid-nitrogen-ground tissue using TRIzol reagent. cDNA was synthesized using the Solg RT kit according to the manufacturer’s instructions. RT-qPCR was performed using Solg 2× Multiplex Real-Time PCR Smart Mix containing SYBR Green on a CFX Duet Real-Time PCR system (Bio-Rad). Cycling conditions were 95°C for 1 min followed by 40 cycles of 95°C for 15 s, 60°C for 15 s, and 72°C for 20 s. Relative transcript abundance was calculated using the 2−ΔΔCt method with ACTIN2 as the reference gene. Primer sequences are provided in Supplementary Table 1.

### Measurement of S-nitrosothiols

*S-*nitrosothiol (SNO) concentrations were measured using a Sievers NOA-280i Nitric Oxide Analyzer (Estero, FL, USA) according to Hussain et al. (2018).

### Electrolyte leakage and oxidative-stress measurements

Electrolyte leakage from Arabidopsis rosette leaves was measured using a conductivity meter (HORIBA Twin Model B-173, Japan) as described by (Rahman et al., 2022). Hydrogen peroxide (H_2_O_2_) accumulation was visualized by 3,3′-diaminobenzidine (DAB) staining according to Das et al. (2022). H_2_O_2_ and malondialdehyde (MDA) contents were quantified as described by Tomassi et al. (2025) and Kim et al. (2020), respectively.

### Enzymatic antioxidant assays

For antioxidant enzyme assays, protein was extracted in phosphate-buffered saline and quantified using the Bradford assay. Catalase, ascorbate peroxidase, and peroxidase activities were quantified according to Rahman et al. (2019).

### Assessment of ABA sensitivity and ABA content

ABA sensitivity assays were performed according to Zhu et al. (2020). Sterilized seeds were grown on half-strength MS medium under a 16-h-light/8-h-dark photoperiod for 4-5 d and then transferred to medium with or without ABA. Plants were photographed 7 d after transfer. Endogenous ABA was quantified according to (Woo et al., 2025), using 100 ng of labeled (±)-3,5,5,7,7,7-d6-ABA as an internal standard.

### Statistical analysis

All statistical analyses, including the Student’s two-tailed *t*-test and analysis of variance (ANOVA) with Tukey’s test, were performed using “ggplot2”, “ggpubr”, “dplyr”, “agricolae”, and “tibble” in RStudio (R 4.4.3). Biological replicates were indicated in the Fig. legends.

## Acknowledgments

We are grateful to the Global Korea Scholarship (GKS) of the Government of the Republic of Korea for its support of A.K.D. We acknowledge the help from previous lab members, Dr. Qari Muhammad Imran and Dr. Noreen Falak, in generating the double mutant.

## Author contributions

A.K.D. and B.-W.Y. conceptualized and designed the study; A.K.D. performed the experimental work and analyzed the data. A.K.D. and M. G.M. wrote the manuscript; D.-S.L., R.A.M., R.A., N.J.M., Y-J.W., S.S., and C-W.P. helped in data curation and analysis; A.H. supervised and helped in experimental setup. S.S. reviewed the manuscript; B.-G.M. and I.-J.L. contributed to the resources. M.G.M., S.K., and B.-W.Y. supervised, edited, and contributed to the resources and funding. All authors have read and approved the contents of this paper.

## Funding

This research was supported by the Basic Science Research Program through the National Research Foundation of Korea (NRF), funded by the Ministry of Education (RS-2023-00245922) and Korea Basic Science Institute (National Research Facilities and Equipment Center, 2021R1A6C101A416) funded by the Ministry of Education, a research grant from the 2026 National University Development Project funded by the Ministry of Education, Republic of Korea. This research was also supported by the Glocal 30 program of Kyungpook National University, funded by the Ministry of Education (MOE).

## Data availability

The data that support the findings of this study are available from the corresponding author upon reasonable request.

## Conflicts of interest

The authors declare no conflicts of interest.

## Notes

### Competing Interest Statement

The authors have declared no competing interest.

## References

Akter S, Perri M, Lavilla-Puerta M, Lichtenauer S, He Y, Shukla V, Carbonare LD, Telara Y, Zhang D, Ferretti B, Gunawardana DM, Myers WK, Barreto P, Giuntoli B, Schwarzländer M, Flashman E, Licausi F (2026) H2O2 repurposes plant O2 sensing to regulate post-hypoxia responses. Nature 653: 1130–1138

Ali MA, Abbas A, Kreil DP, Bohlmann H (2013) Overexpression of the transcription factor RAP2.6 leads to enhanced callose deposition in syncytia and enhanced resistance against the beet cyst nematode Heterodera schachtiiin Arabidopsis roots. BMC Plant Biology 13: 47

Arnon DI (1949) Copper Enzymes in Isolated Chloroplasts. Polyphenoloxidase in Beta Vulgaris. Plant Physiology 24: 1–15

Bailey-Serres J, Voesenek LACJ (2008) Flooding Stress: Acclimations and Genetic Diversity. Annual Review of Plant Biology 59: 313–339

Bogos B, Ughy B, Domonkos I, Laczkó-Dobos H, Komenda J, Abasova L, Cser K, Vass I, Sallai A, Wada H, Gombos Z (2010) Phosphatidylglycerol depletion affects photosystem II activity in Synechococcus sp. PCC 7942 cells. Photosynthesis Research 103: 19–30

Chen L, Liao B, Qi H, Xie L-J, Huang L, Tan W-J, Zhai N, Yuan L-B, Zhou Y, Yu L-J, Chen Q-F, Shu W, Xiao S (2015) Autophagy contributes to regulation of the hypoxia response during submergence in Arabidopsis thaliana. Autophagy 11: 2233–2246

Das AK, Anik TR, Rahman MM, Keya SS, Islam MR, Rahman MA, Sultana S, Ghosh PK, Khan S, Ahamed T, Ghosh TK, Tran LS-P, Mostofa MG (2022) Ethanol Treatment Enhances Physiological and Biochemical Responses to Mitigate Saline Toxicity in Soybean. Plants 11: 272

Das AK, Khan I, Yun B-W, Mostofa MG (2026) Beyond hypoxia: Context-dependent transcriptional reprogramming by ERFVIIs. Journal of Integrative Plant Biology

Das AK, Lee D-S, Lee G-J, Kim Y-S, Hussain S, Lee M-S, Yun B-W, Mun B-G (2025) The Central Role of GSNOR: Decoding Nitric Oxide Signaling for Crop Stress Tolerance. International Journal of Molecular Sciences 26: 11486

Das AK, Mostofa MG, Lee D-S, Yun B-W (2026) AP2/ERF transcription factor RAP2.6 regulates early flowering in Arabidopsis thaliana by altering S-nitrosothiol levels and cytokinin responses. bioRxiv: 725052

Dey S, Corina Vlot A (2015) Ethylene responsive factors in the orchestration of stress responses in monocotyledonous plants. Frontiers in Plant Science 6: 640

Dichio C, Sofo A, Yaghoubi Khanghahi M, Crecchio C, Addesso R (2025) Physiological and Biochemical Mechanisms of Waterlogging Tolerance in Plants. Physiologia Plantarum 177: e70663

DuBois M, Gilles KA, Hamilton JK, Rebers PA, Smith F (1956) Colorimetric method for determination of sugars and related substances. Analytical chemistry 28: 350–356

Falak N, Imran QM, Hussain A, Yun B-W (2021) Transcription Factors as the “Blitzkrieg” of Plant Defense: A Pragmatic View of Nitric Oxide’s Role in Gene Regulation. International Journal of Molecular Sciences 22: 522

Fowler S, Thomashow MF (2002) Arabidopsis Transcriptome Profiling Indicates That Multiple Regulatory Pathways Are Activated during Cold Acclimation in Addition to the CBF Cold Response Pathway. The Plant Cell 14: 1675–1690

Gibbs DJ, Conde JV, Berckhan S, Prasad G, Mendiondo GM, Holdsworth MJ (2015) Group VII Ethylene Response Factors Coordinate Oxygen and Nitric Oxide Signal Transduction and Stress Responses in Plants. Plant Physiology 169: 23–31

Giuntoli B, Perata P (2017) Group VII Ethylene Response Factors in Arabidopsis: Regulation and Physiological Roles Plant Physiology 176: 1143–1155

Guo F-Q, Okamoto M, Crawford NM (2003) Identification of a Plant Nitric Oxide Synthase Gene Involved in Hormonal Signaling. Science 302: 100–103

Gupta KJ, Igamberdiev AU (2016) Reactive Nitrogen Species in Mitochondria and Their Implications in Plant Energy Status and Hypoxic Stress Tolerance. Frontiers in Plant Science 7: 369

Hartman S, Liu Z, Van Veen H, Vicente J, Reinen E, Martopawiro S, Zhang H, Van Dongen N, Bosman F, Bassel GW (2019) Ethylene-mediated nitric oxide depletion pre-adapts plants to hypoxia stress. Nature Communications 10: 4020

Hartman S, Sasidharan R, Voesenek LACJ (2021) The role of ethylene in metabolic acclimations to low oxygen. New Phytologist 229: 64–70

He Y, Tang R-H, Hao Y, Stevens RD, Cook CW, Ahn SM, Jing L, Yang Z, Chen L, Guo F, Fiorani F, Jackson RB, Crawford NM, Pei Z-M (2004) Nitric Oxide Represses the Arabidopsis Floral Transition. Science 305: 1968–1971

Hess N, Klode M, Anders M, Sauter M (2011) The hypoxia responsive transcription factor genes ERF71/HRE2 and ERF73/HRE1 of Arabidopsis are differentially regulated by ethylene. Physiologia Plantarum 143: 41–49

Hussain A, Mun B-G, Imran QM, Lee S-U, Adamu TA, Shahid M, Kim K-M, Yun B-W (2016) Nitric Oxide Mediated Transcriptome Profiling Reveals Activation of Multiple Regulatory Pathways in Arabidopsis thaliana. Frontiers in Plant Science Volume 7 - 2016

Hussain A, Yun B-W, Loake GJ (2018) Nitric Oxide Analyzer Quantification of Plant S-Nitrosothiols. In A Mengel, C Lindermayr, eds, Nitric Oxide: Methods and Protocols. Springer New York, New York, NY, pp 223–230

Jisha V, Dampanaboina L, Vadassery J, Mithöfer A, Kappara S, Ramanan R (2015) Overexpression of an AP2/ERF Type Transcription Factor OsEREBP1 Confers Biotic and Abiotic Stress Tolerance in Rice. PLOS ONE 10: e0127831

Kim TY, Ku H, Lee S-Y (2020) Crop Enhancement of Cucumber Plants under Heat Stress by Shungite Carbon. International Journal of Molecular Sciences 21: 4858

Krishnaswamy S, Verma S, Rahman MH, Kav NNV (2011) Functional characterization of four APETALA2-family genes (RAP2.6, RAP2.6L, DREB19 and DREB26) in Arabidopsis. Plant Molecular Biology 75: 107–127

Liao K, Wang L-N, Lu X-Y, Zhang H-M, Yan Z-R, Zhang T-Y, Li Y, Xu L-J, Zhong Q-L, Chen Q-F, Zhou Y, Xie R-H, Xiao S (2026) The SnRK1–RAP2.4h–PIP2 module contributes to the trade-off between growth and hypoxia tolerance in plants. Nature Communications 17: 5886

Licausi F, Kosmacz M, Weits DA, Giuntoli B, Giorgi FM, Voesenek LACJ, Perata P, van Dongen JT (2011) Oxygen sensing in plants is mediated by an N-end rule pathway for protein destabilization. Nature 479: 419–422

Lichtenthaler HK (1987) Chlorophylls and carotenoids: Pigments of photosynthetic biomembranes. *In* Methods in Enzymology, Vol 148. Academic Press, pp 350–382

Lichtenthaler HK, Wellburn AR (1983) Determinations of total carotenoids and chlorophylls a and b of leaf extracts in different solvents. Biochemical Society Transactions 11: 591–592

Liu M, Wei J-W, Liu W, Gong B (2023) S-nitrosylation of ACO homolog 4 improves ethylene synthesis and salt tolerance in tomato. New Phytologist 239: 159–173

Liu Z, Hartman S, van Veen H, Zhang H, Leeggangers HACF, Martopawiro S, Bosman F, de Deugd F, Su P, Hummel M, Rankenberg T, Hassall KL, Bailey-Serres J, Theodoulou FL, Voesenek LACJ, Sasidharan R (2022) Ethylene augments root hypoxia tolerance via growth cessation and reactive oxygen species amelioration. Plant Physiology 190: 1365–1383

Loreti E, Perata P (2023) ERFVII transcription factors and their role in the adaptation to hypoxia in Arabidopsis and crops. Frontiers in Genetics Volume 14 - 2023

Luan H, Guo B, Shen H, Pan Y, Hong Y, Lv C, Xu R (2020) Overexpression of Barley Transcription Factor HvERF2.11 in Arabidopsis Enhances Plant Waterlogging Tolerance. International Journal of Molecular Sciences 21: 1982

Lv Y, Fu S, Chen S, Zhang W, Qi C (2016) Ethylene response factor BnERF2-like (ERF2.4) from Brassica napus L. enhances submergence tolerance and alleviates oxidative damage caused by submergence in Arabidopsis thaliana. The Crop Journal 4: 199–211

Nakano T, Suzuki K, Fujimura T, Shinshi H (2006) Genome-Wide Analysis of the ERF Gene Family in Arabidopsis and Rice. Plant Physiology 140: 411–432

Praveen A, Singh S (2026) Flooding stress resilience and crop improvement. Plant Cell Reports 45: 264

Rahman MM, Mostofa MG, Das AK, Anik TR, Keya SS, Ahsan SM, Khan MAR, Ahmed M, Rahman MA, Hossain MM, Tran L-SP (2022) Ethanol Positively Modulates Photosynthetic Traits, Antioxidant Defense and Osmoprotectant Levels to Enhance Drought Acclimatization in Soybean. Antioxidants 11: 516

Rahman MM, Mostofa MG, Rahman MA, Islam MR, Keya SS, Das AK, Miah MG, Kawser AQMR, Ahsan SM, Hashem A, Tabassum B, Abd_Allah EF, Tran L-SP (2019) Acetic acid: a cost-effective agent for mitigation of seawater-induced salt toxicity in mung bean. Scientific Reports 9: 15186

Sakurai I, Hagio M, Gombos Z, Tyystja□rvi T, Paakkarinen V, Aro E-M, Wada H (2003) Requirement of Phosphatidylglycerol for Maintenance of Photosynthetic Machinery. Plant Physiology 133: 1376–1384

Samant SB, Swain J, Yadav N, Yadav R, Singh P, Rai P, Sheri V, Sreeman S, Subramanyam R, Pareek A, Gupta KJ (2025) Overexpression of Phytoglobin1 in Rice Leads to Enhanced Nitrogen Use Efficiency via Modulation of Nitric Oxide. Plant, Cell & Environment 48: 2755–2768

Sasidharan R, Hartman S, Liu Z, Martopawiro S, Sajeev N, van Veen H, Yeung E, Voesenek LACJ (2017) Signal Dynamics and Interactions during Flooding Stress. Plant Physiology 176: 1106–1117

Sherwood OL, Carroll R, Burke S, McCabe PF, Kacprzyk J (2023) A simple and cost-effective method for studying anoxia tolerance in plants. Applications in Plant Sciences 11: e11509

Somogyi M (1952) NOTES ON SUGAR DETERMINATION. Journal of Biological Chemistry 195: 19–23

Tomassi AH, Perea-García A, Rodrigo G, Sánchez-Vicente J, Cisneros AE, Olmos M, Andrés-Bordería A, López-Dolz L, Daròs J-A, Peñarrubia L, Carbonell A (2025) Arabidopsis thaliana iron superoxide dismutase FeSOD1 protects ARGONAUTE 1 in a copper-dependent manner. Journal of Experimental Botany 76: 5465–5480

van der Wijk E, Xu J, Sasidharan R (2026) Phytoglobin 1 in plant hypoxia acclimation and development: integrating oxygen sensing, nitric oxide homeostasis, and redox balance. Journal of Experimental Botany: erag147

Wang L-N, Wang W-C, Liao K, Xu L-J, Xie D-X, Xie R-H, Xiao S (2025) Survival mechanisms of plants under hypoxic stress: Physiological acclimation and molecular regulation. Journal of Integrative Plant Biology 67: 440–454

Wang Z, Cao G, Wang X, Miao J, Liu X, Chen Z, Qu L-J, Gu H (2008) Identification and characterization of COI1-dependent transcription factor genes involved in JA-mediated response to wounding in Arabidopsis plants. Plant Cell Reports 27: 125–135

White MD, Klecker M, Hopkinson RJ, Weits DA, Mueller C, Naumann C, O’Neill R, Wickens J, Yang J, Brooks-Bartlett JC, Garman EF, Grossmann TN, Dissmeyer N, Flashman E (2017) Plant cysteine oxidases are dioxygenases that directly enable arginyl transferase-catalysed arginylation of N-end rule targets. Nature Communications 8: 14690

Woo J-I, Adhikari A, Gam H-J, Jeon JR, Lee D-S, Kwon E-H, Kang S-M, Yun B-W, Lee I-J (2025) Integrated role of biochar and PGPR (Leclercia adecarboxylata HW04) in enhancing cadmium phytoremediation and stress tolerance in Glycine max L. Plant Physiology and Biochemistry 220: 109489

Xuan S, Cheng M, Wen Y, Wei Q, Zhang D, He P, Yu J (2026) ABI5-activated ALKBH10B demethylates RAP2.6 mRNA to modulate Arabidopsis salt tolerance. Plant Cell Reports 45: 234

Yamaoka Y, Shin S, Lee Y, Ito M, Lee Y, Nishida I (2021) Phosphatidylserine Is Required for the Normal Progression of Cell Plate Formation in Arabidopsis Root Meristems. Plant and Cell Physiology 62: 1396–1408

Yao S, Kim S-C, Li J, Tang S, Wang X (2024) Phosphatidic acid signaling and function in nuclei. Progress in Lipid Research 93: 101267

Yin D, Sun D, Han Z, Ni D, Norris A, Jiang C-Z (2019) PhERF2, an ethylene-responsive element binding factor, plays an essential role in waterlogging tolerance of petunia. Horticulture Research 6: 83

Zhan N, Wang C, Chen L, Yang H, Feng J, Gong X, Ren B, Wu R, Mu J, Li Y, Liu Z, Zhou Y, Peng J, Wang K, Huang X, Xiao S, Zuo J (2018) S-Nitrosylation Targets GSNO Reductase for Selective Autophagy during Hypoxia Responses in Plants. Molecular Cell 71: 142–154.e146

Zhang Y, Feng Y, Liu Y, Qiu Y, Zhang X, Cheng J, Wang Y (2026) GSNOR mediates NO signaling to enhance salt tolerance in tobacco by regulating redox and photosynthesis. Photosynthesis Research 164: 21

Zhu Q, Zhang J, Gao X, Tong J, Xiao L, Li W, Zhang H (2010) The Arabidopsis AP2/ERF transcription factor RAP2.6 participates in ABA, salt and osmotic stress responses. Gene 457: 1–12

Zhu Y, Huang P, Guo P, Chong L, Yu G, Sun X, Hu T, Li Y, Hsu C-C, Tang K, Zhou Y, Zhao C, Gao W, Tao WA, Mengiste T, Zhu J-K (2020) CDK8 is associated with RAP2.6 and SnRK2.6 and positively modulates abscisic acid signaling and drought response in Arabidopsis. New Phytologist 228: 1573–1590

