## Supplementary File for "RAP2.6 negatively regulates Arabidopsis submergence tolerance through GSNOR1-dependent redox and hormonal signaling"

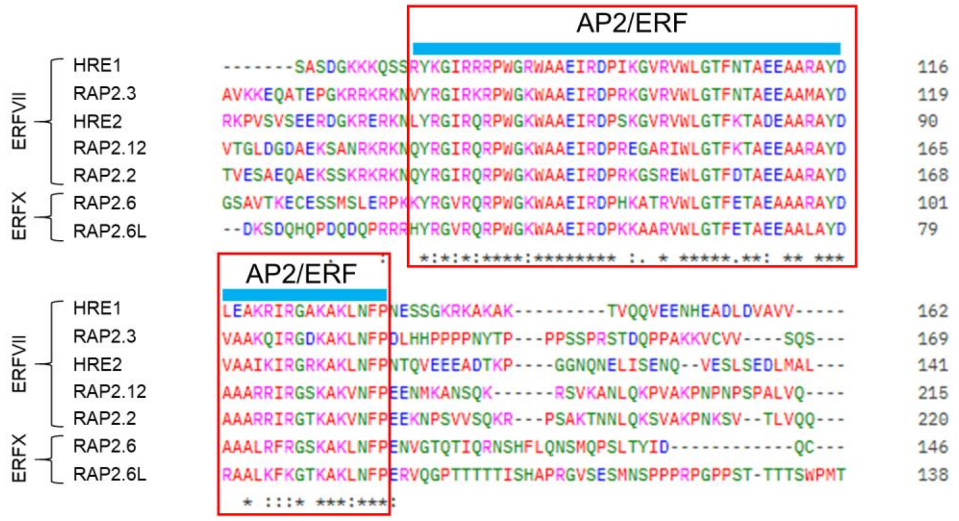


**Fig. S1.** Multiple sequence alignment of AP2/ERF transcription factors (ERFVII and ERFX) using CLUSTAL 0 (1.2.4).

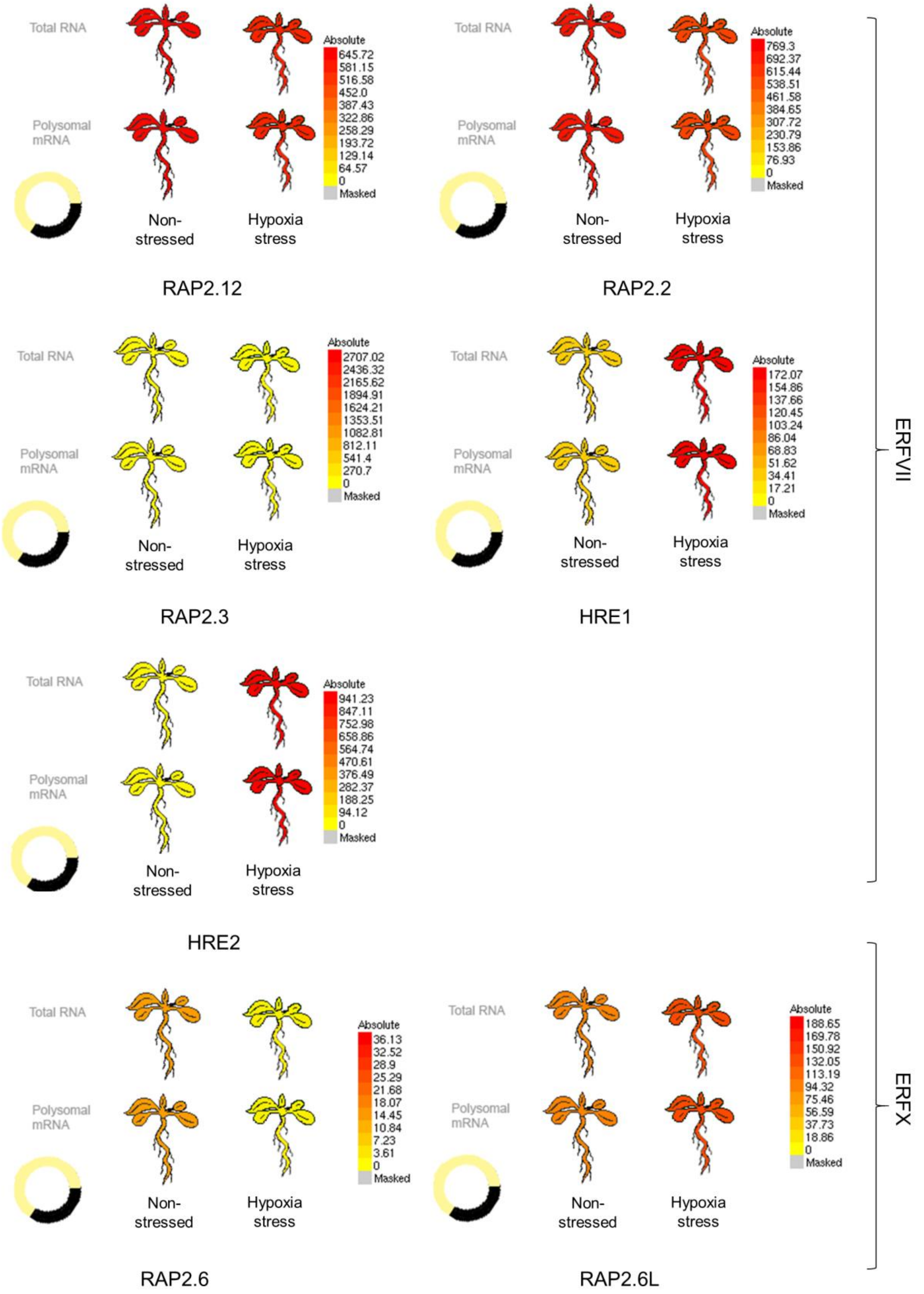


**Fig. S2.** Visualization of the expression levels of AP2/ERF transcription under hypoxia, retrieved from Arabidopsis eFP Browser.

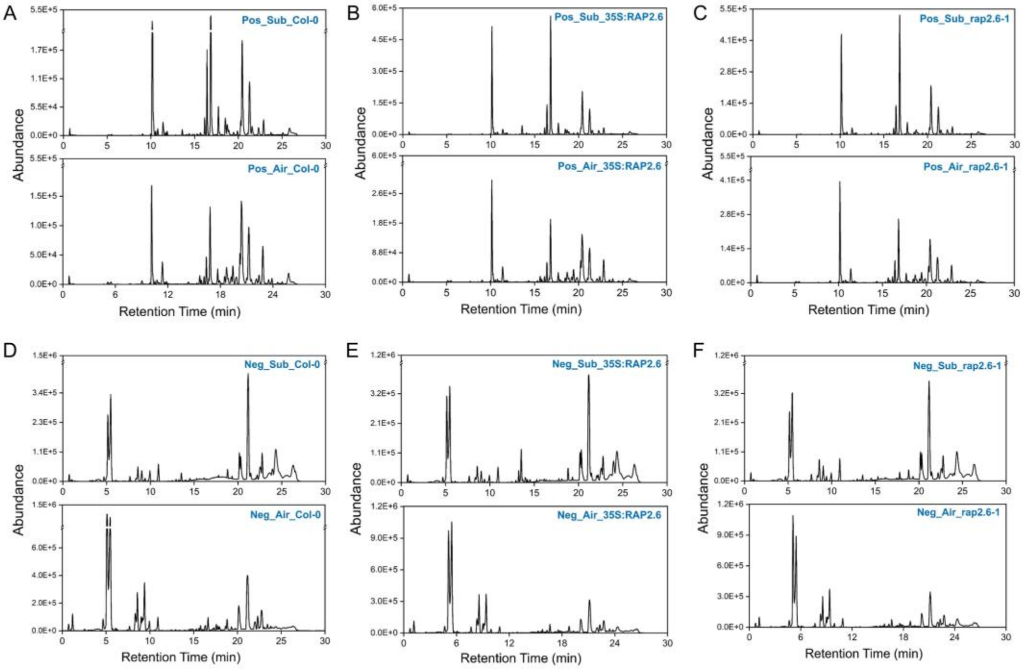


**Fig. S3.** LC Chromatogram showing the metabolite differences based on abundance of control and treatment samples – Col-0, 35S::RAP2.6, and rap2.6-1 in (A–C) $[M+H]^+$ ion mode and (D–F) $[M-H]^-$ ion mode, after 1 day of dark submergence or air.

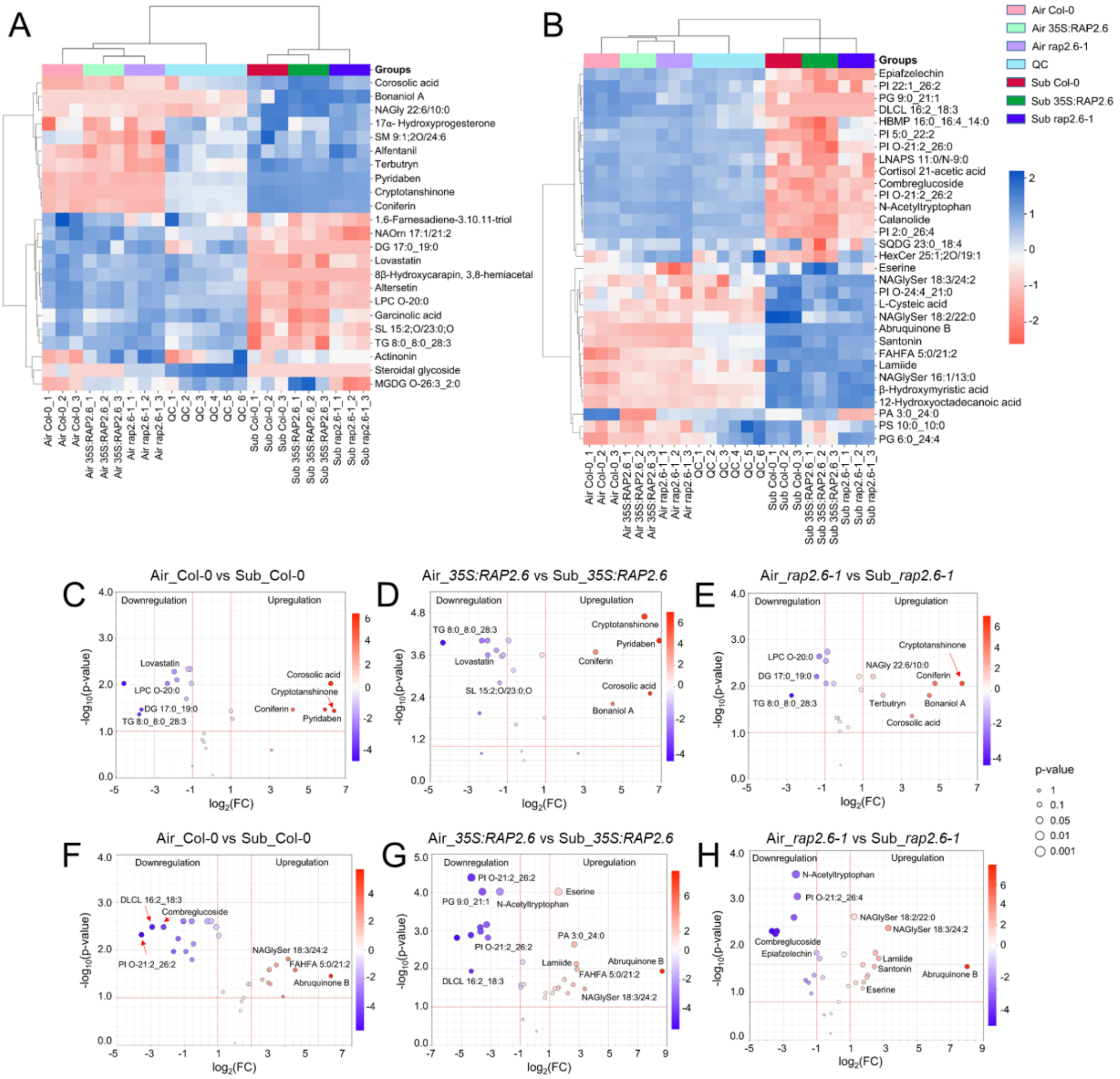


**Fig. S4.** Metabolic profiling of RAP2.6 under submergence stress. Dendrograms show hierarchical relationships in Col-0, *35S::RAP2.6*, and *rap2.6-1* plants after 1 day of dark submergence or air, in (A) $[M+H]^+$ and (B) $[M-H]^-$ ion modes. Volcano plot showing the statistical significance of metabolite changes between air and submergence treatment in (C) Col-0, (D) *35S::RAP2.6*, and (E) *rap2.6-1* plants in $[M+H]^+$ ion mode, and in (F) Col-0, (G) *35S::RAP2.6*, and (H) *rap2.6-1* plants in $[M-H]^-$ ion mode. Each dot represents one detected metabolite, plotted according to $\log_2$ fold change (FC) with respect to $-\log_{10}$(p-value). Positive $\log_2$FC values indicate upregulated metabolites in the treatment group, whereas negative values indicate downregulated metabolites relative to air treatment. The color of the dots represents the direction and magnitude of change from downregulated (purple) to upregulated (red) features, and the point size scales with statistical significance.

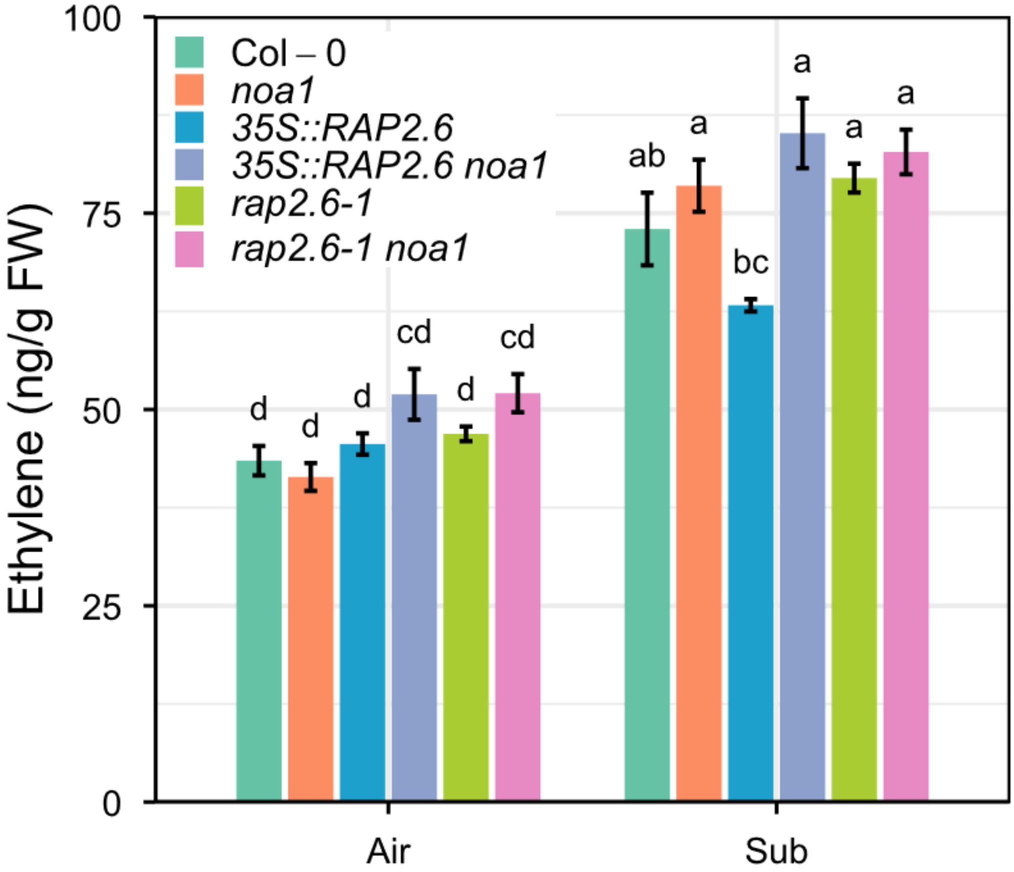


**Fig. S5. (**A) Ethylene levels in Col-0, *noa1*, *35S::RAP2.6*, *35S::RAP2.6 noa1*, *rap2.6-1*, and *rap2.6-1 noa1* plants exposed to air or dark submergence for 1 day. Bars represent means ± SE (*n* = 3) of three independent experiments (for one experiment, 4 plants were used for each genotype). Different letters indicate statistically significant differences by two-way ANOVA and Tukey's HSD ($P < 0.05$).

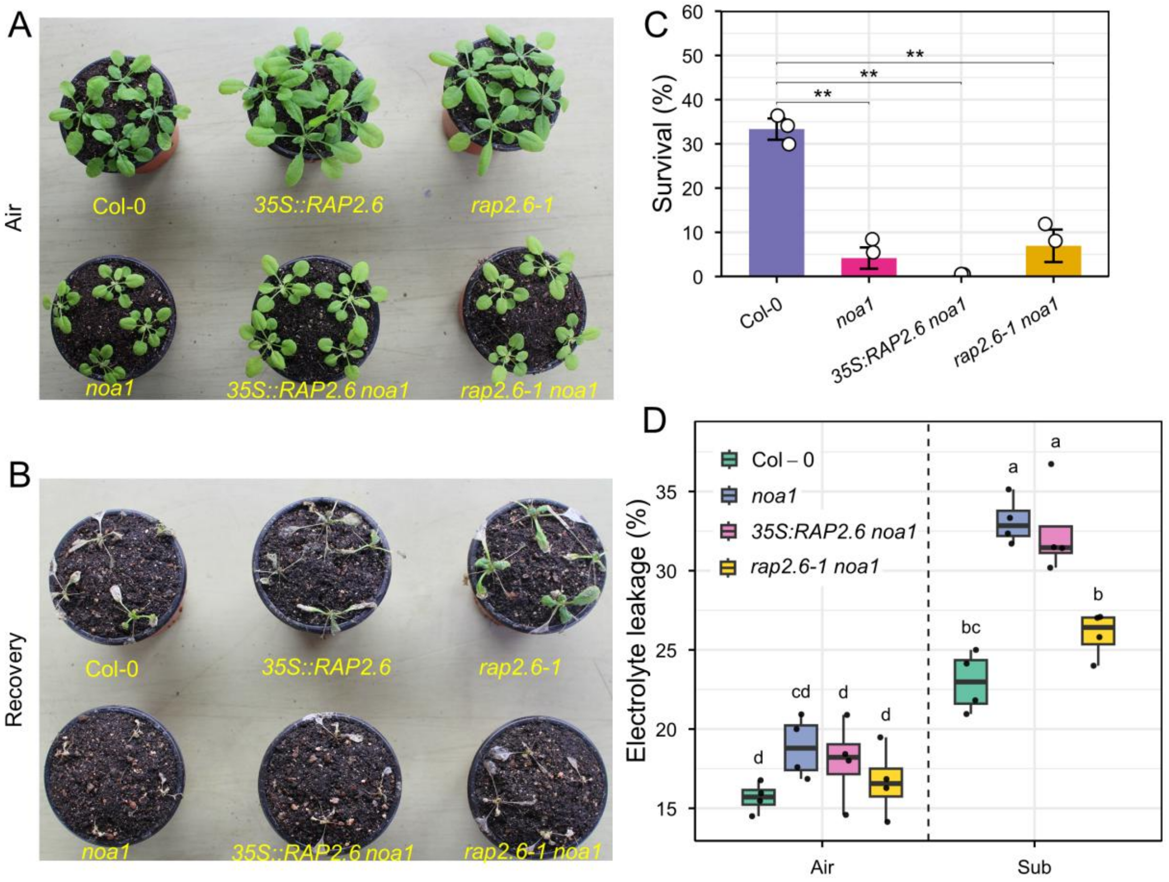


**Fig. S6.** (A, B) Phenotypes of Col-0, *35S::RAP2.6*, *rap2.6-1*, *noa1*, *35S::RAP2.6 noa1*, and *rap2.6-1 noa1* plants were photographed before submergence (air) and after 5 days of dark submergence, followed by 7 days of recovery. (C) Survival rates were calculated after 7 days of recovery. Bars represent means ± SE ($n$ = 3) of three independent experiments (for one experiment, >10 plants were scored for each genotype). (D) Electrolyte leakage in Col-0, *noa1*, *35S::RAP2.6 noa1*, and *rap2.6-1 noa1* plants exposed to air or 2 days of dark submergence (means ± SE, $n$ = 4 biological replicates for EL, $n$ = 9 technical replicates from 3 biological replicates). Asterisks indicate significant differences by Student's two-tailed *t*-test ($*P < 0.05$, $**P < 0.01$, $***P < 0.001$, $****P < 0.0001$). Different letters indicate statistically significant differences by two-way ANOVA and Tukey's HSD ($P < 0.05$).

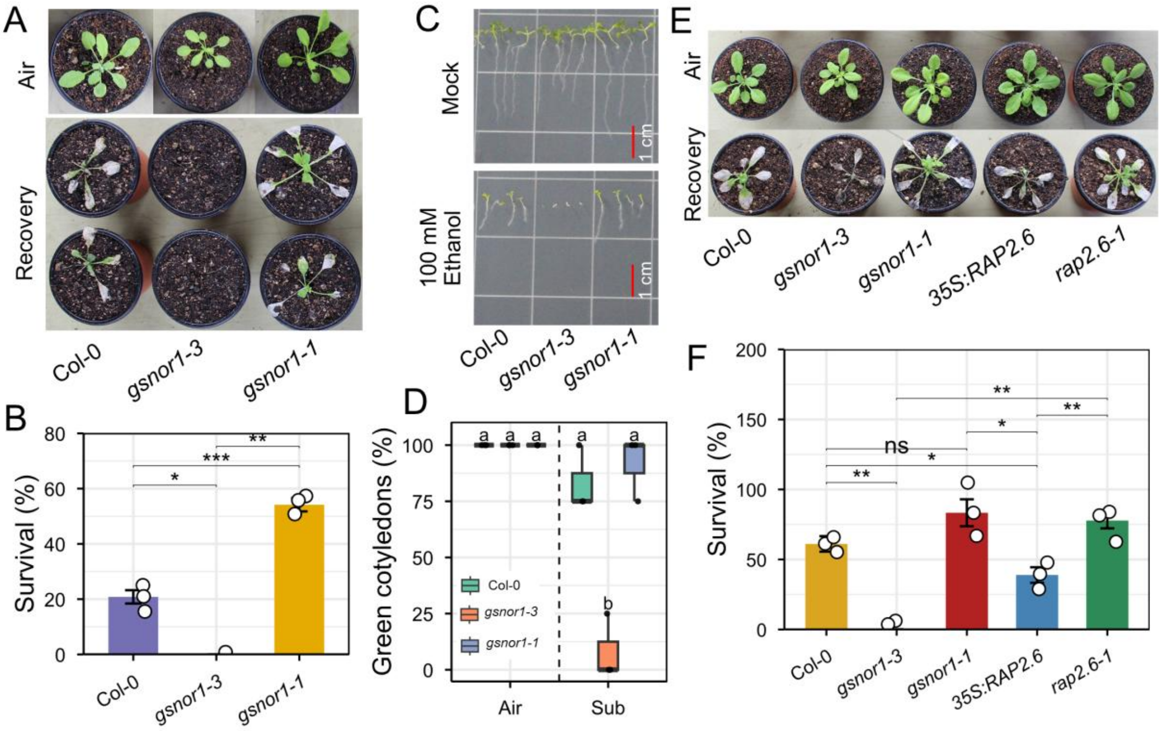


**Fig. S7.** (A) Phenotypes of Col-0, *gsnor1-3*, and *gsnor1-1* plants were photographed before submergence (air) and after 5 days of dark submergence, followed by 7 days of recovery. (B) Survival rates were calculated after 7 days of recovery. Bars represent means ± SE ($n$ = 3) of three independent experiments (for one experiment, >10 plants were scored for each genotype). (C) Phenotypes and (D) green cotyledon percentage of Col-0, *gsnor1-3*, and *gsnor1-1* seedlings grown on ½ MS media for 7 days, with or without 100 mM Ethanol. Bars represent means ± SE ($n$ = 3 biological replicates) of three independent experiments (for one experiment, 4 plants were scored for each genotype). (E) Phenotypes of Col-0, *gsnor1-3*, *gsnor1-1*, *35S::RAP2.6*, and *rap2.6-1* plants were photographed before submergence (air) and after 5 days of dark submergence, followed by 7 days of recovery. (F) Survival rates were calculated after 7 days of recovery. Bars represent means ± SE ($n$ = 3) of three independent experiments (for one experiment, >10 plants were scored for each genotype). Asterisks indicate significant differences by Student's two-tailed $t$-test ($*P < 0.05$, $**P < 0.01$, $***P < 0.001$, $****P < 0.0001$). Different letters indicate statistically significant differences by two-way ANOVA and Tukey's HSD ($P < 0.05$).

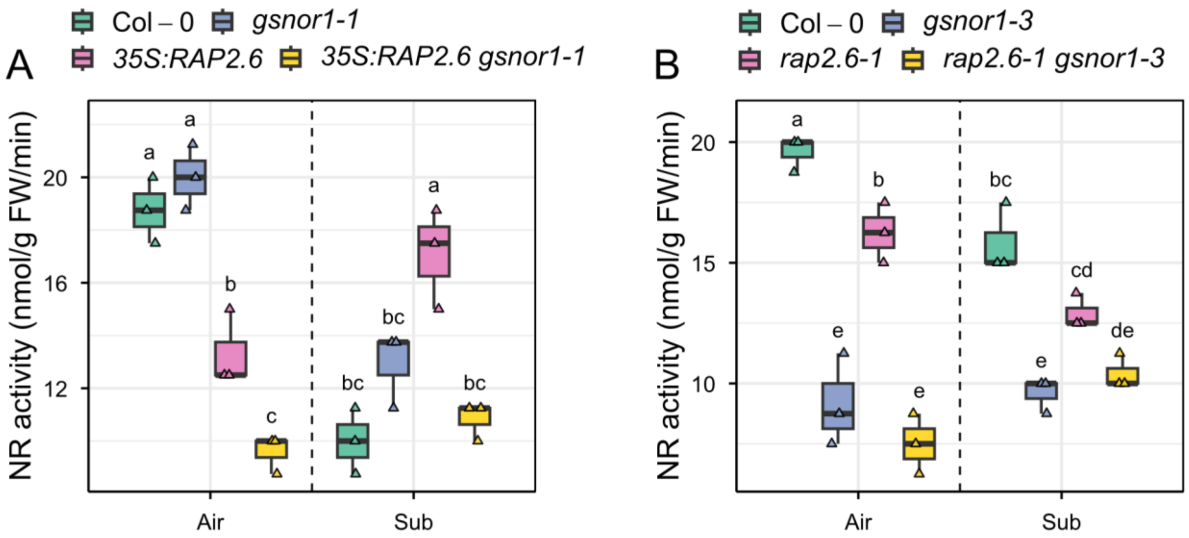


**Fig. S8.** (A, B) Nitrate reductase activity in Col-0, *gsnor1-1*, *35S::RAP2.6*, and *35S::RAP2.6 gsnor1-1*, *and* Col-0, *gsnor1-3*, *rap2.6-1*, and *rap2.6-1 gsnor1-3* plants exposed to air or dark submergence for 1 day (means ± SE, $n$ = 3 biological replicates). Different letters indicate statistically significant differences by two-way ANOVA and Tukey's HSD ($P < 0.05$).

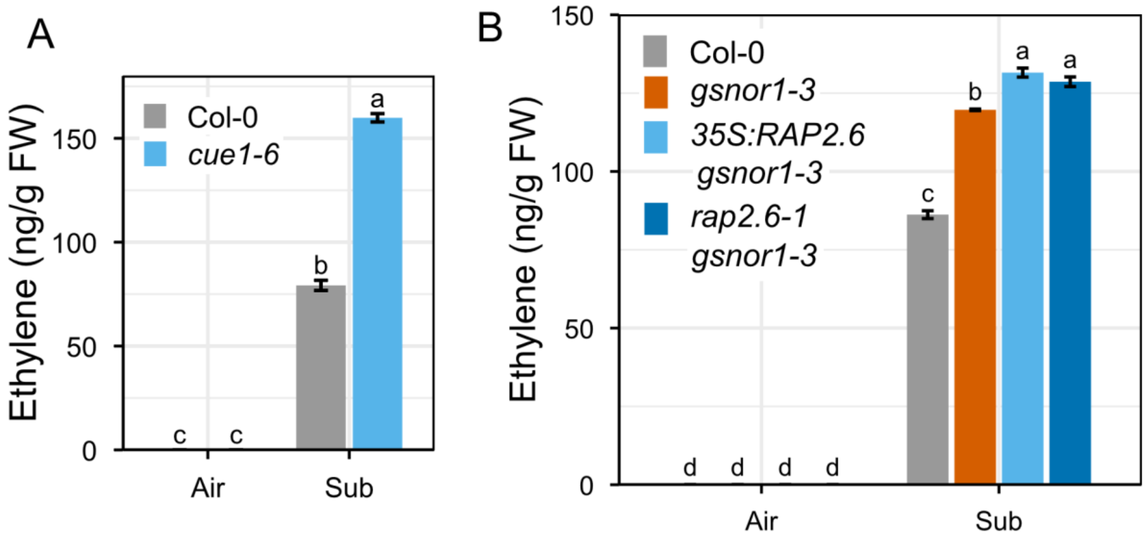


**Fig. S9.** (A) Ethylene levels in Col-0 and *cue1-6* plants exposed to air or dark submergence for 1 day. (B) Ethylene levels in Col-0, *gsnor1-3*, *35S::RAP2.6 gsnor1-3*, and *rap2.6-1 gsnor1-3* plants exposed to air or dark submergence for 1 day. Bars represent means ± SE ($n$ = 3) of three independent experiments (for one experiment, 4 plants were used for each genotype). Different letters indicate statistically significant differences by two-way ANOVA and Tukey's HSD ($P < 0.05$).

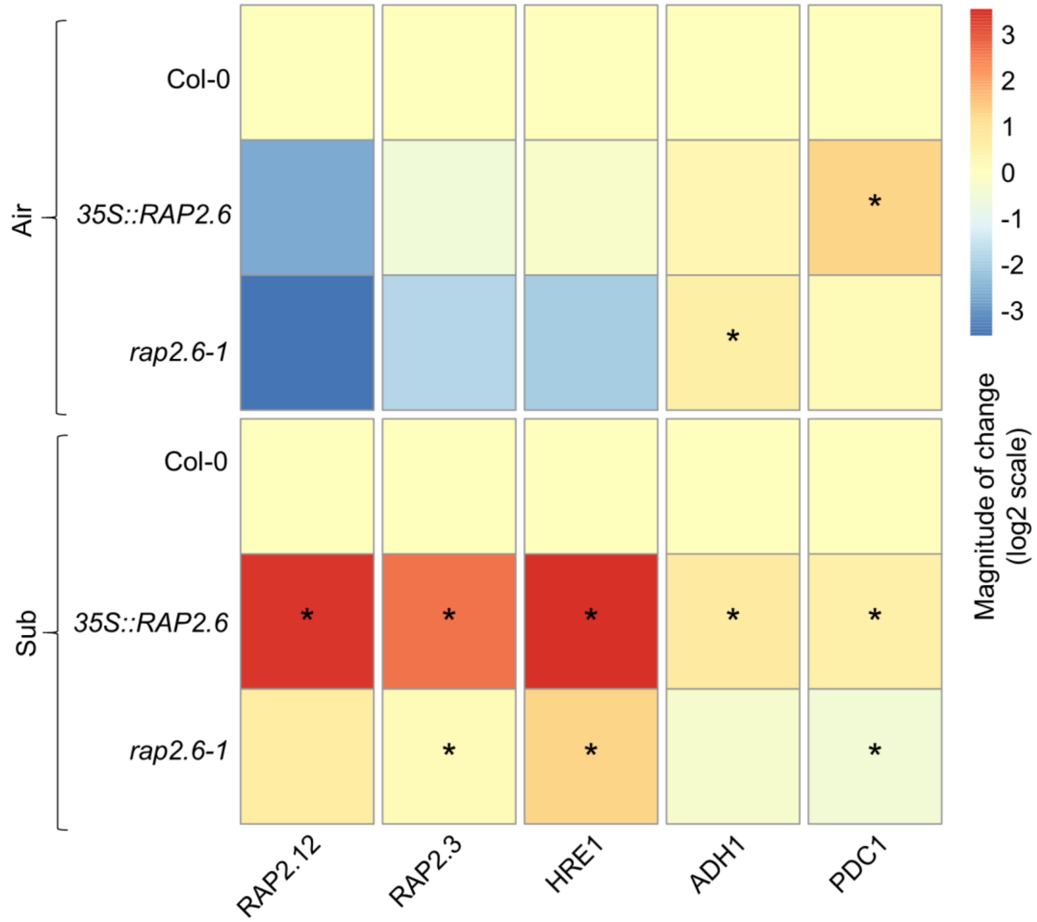


**Fig. S10.** Heatmap showing the relative gene expression of *RAP2.12*, *RAP2.3*, *HRE1*, *ADH1*, and *PDC1* in Col-0, *35S::RAP2.6*, and *rap2.6-1* plant leaves exposed to air or dark submergence for 24 hours (means ± SE, $n = 3$ biological replicates). Asterisks indicate significant differences by Student's two-tailed $t$-test ($*P < 0.05$, $**P < 0.01$, $***P < 0.001$, $****P < 0.0001$).

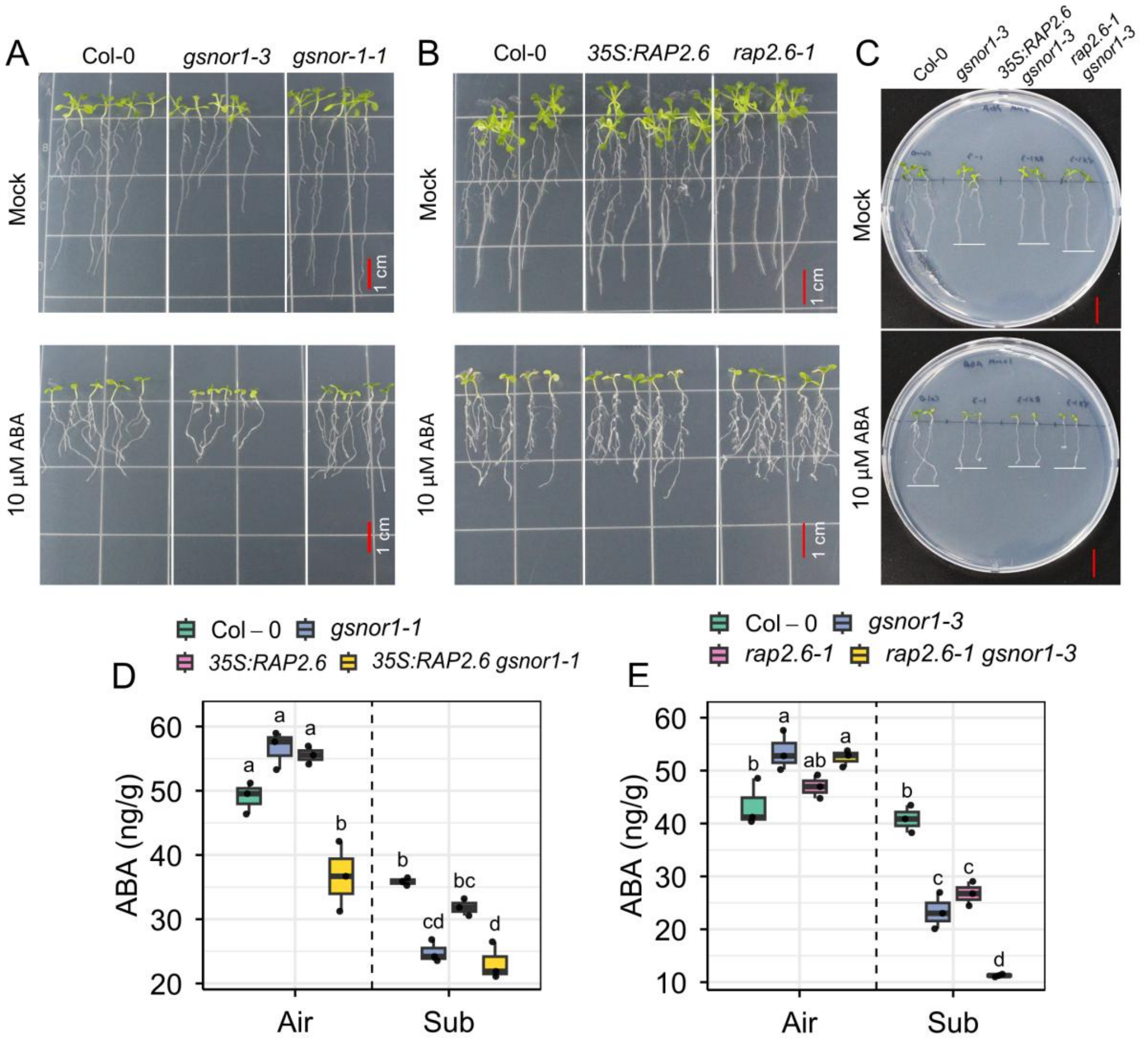


**Fig. S11.** (A, B) Phenotypes of Col-0, *gsnor1-3*, and *gsnor1-1*, and Col-0, *35S::RAP2.6*, and *rap2.6-1* seedlings grown on ½ MS media for 7 days, with or without 10 µM ABA. (C) Phenotypes of Col-0, *gsnor1-3*, *35S::RAP2.6 gsnor1-1*, and *35S::RAP2.6 gsnor1-3* seedlings grown on ½ MS media for 7 days, with or without 10 µM ABA. (D, E) ABA level in Col-0, *gsnor1-1*, *35S::RAP2.6*, and *35S::RAP2.6 gsnor1-1, and* Col-0, *gsnor1-3*, *rap2.6-1*, and *rap2.6-1 gsnor1-3* plants exposed to air or dark submergence for 1 day (means ± SE, $n$ = 3 biological replicates). Different letters indicate statistically significant differences by two-way ANOVA and Tukey's HSD ($P < 0.05$).

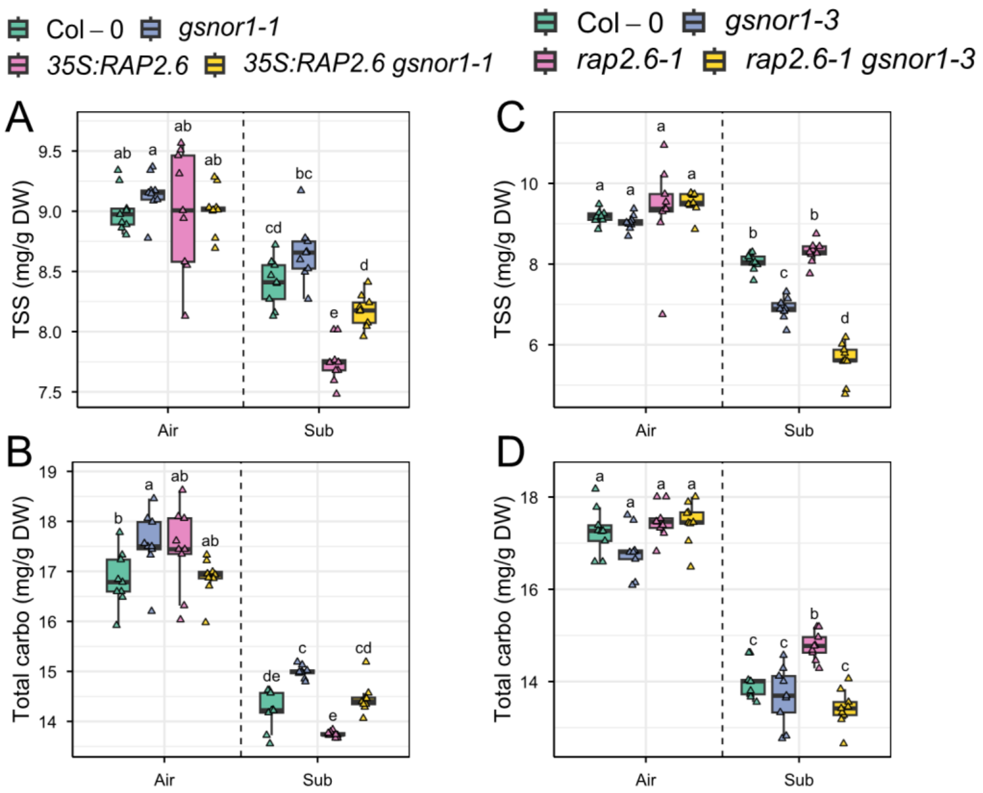


**Fig. S12.** (A) Total soluble sugar (TSS) and (B) total carbohydrate (carbo) in Col-0, *gsnor1-1*, *35S::RAP2.6*, and *35S::RAP2.6 gsnor1-1* plants exposed to air or dark submergence for 2 days (means ± SE, *n* = 9 technical replicates from 3 biological replicates). (C) Total soluble sugar (TSS) and (D) total carbohydrate (carbo) in Col-0, *gsnor1-3*, *rap2.6-1*, and *rap2.6-1 gsnor1-3* plants exposed to air or dark submergence for 2 days (means ± SE, *n* = 9 technical replicates from 3 biological replicates). Different letters indicate statistically significant differences by two-way ANOVA and Tukey's HSD ($P < 0.05$). DW, freeze-dry weight.

**Table S1.** Gene-specific primer lists.

| **For qRT-PCR** | Sequence (5' --> 3') |
|---|---|
| *AtActin2 F* | AGACCAGCTCTTCCATCGAG |
| *AtActin2 R* | GCAGCTTCCATTCCCACAAA |
| *AtRAP2.12 F* | AGCGAGTTTATTTGGCCGGA |
| *AtRAP2.12 R* | AAAACTGAACCTTCCGCAGC |
| *AtRAP2.3 F* | CCGATTATGCCCCTCTCGTC |
| *AtRAP2.3 R* | GGGATGGAGTTTGGAGGTGG |
| *AtHRE1 F* | ACATAGCGCCGGAGAAGATT |
| *AtHRE1 R* | CCATCTGATGCTGAGCCTGAA |
| *AtADH1 F* | AGTGGTTCACTCTGGTCAGG |
| *AtADH1 R* | AAACCAACAGCACCAAGACC |
| *AtPDC1 F* | CCTTCCTCCGCTATCACCAT |
| *AtPDC1 R* | CGGCGTTTAGCTCGTTACAA |
| *AtACO4 F* | ATGTCCCTGATCTCGACGAC |
| *AtACO4 R* | GATTCTCGCACAGCAGATCC |
| *AtACO5 F* | GCCATTACCCTCCTTGTCCT |
| *AtACO5 R* | GCTGAACATCGATCCACTCG |
| *AtPCO1 F* | GGTTTAACTCCGACCATGCC |
| *AtPCO1 R* | AACACCAGAAGGTGGCAAAC |
| *AtPCO2 F* | TTCGCTGATGGCAAATCTGG |
| *AtPCO2 R* | GATCGTCCGGTCACTGTAGA |
| *AtCDK8 F* | AGCCAACCGATGGAGAAGAA |
| *AtCDK8 R* | CATGAGCTCCTGCCATGTTC |
| *AtNCED3 F* | TCTCCGGTGGTTTACGACAA |
| *AtNCED3 R* | GGACCCTATCACGACGACTT |
| *AtSnRK2.6 F* | TGCAGATCATTGCAGAAGCA |
| *AtSnRK2.6 R* | TCCGCTACTGTCGATGTCAA |
| *AtRD29A F* | TGATGGTCAACGGAAGGTCA |
| *AtRD29A R* | ATCTCCGGTACTCCTCCAGT |
| *AtCOR15A F* | GAGAGAAGTCACTGCCTGGT |
| *AtCOR15A R* | CAACTTCTACGCCCTTGACG |
| **For genotyping** | Sequence (5' --> 3') |
| *AtRAP2.6-1_F* (Ali et al., 2013) | TGATGTCGGCAGTTACAAGTG |
| *AtRAP2.6-1_R* (Ali et al., 2013) | TTCCTCTAAAGCGAAGTGCTG |
| *Atgsnor1-1_F* | TATATAATGGTTCGACGCATATTT |
| *Atgsnor1-1_R* | GTCGGGTCGGGTCGTTAATA |
| *Atgsnor1-1_Sail LB* | TTCATAACCAATCTGGATACA |
| *Atgsnor1-3_F* | TATATAATGGTTCGACGCATA |
| *Atgsnor1-3_R* | CCACCAACACTCTCAACAATC |
| *Atgsnor1-3_GABI LB* | ATATTGACCATCATACTCATT |
| *Atnos1(Crawford primer)_F* | CCTGCAACACAACAAGTTGACG |
| *Atnos1(Crawford primer)_R* | GGGCGGATATAACCGTCTCCA |
